# Du-IN-v2: Unleashing the Power of Vector Quantization for Decoding Cognitive States from Intracranial Neural Signals

**DOI:** 10.1101/2024.10.16.618686

**Authors:** Hui Zheng, Hai-Teng Wang, Wei-Bang Jiang, Zhong-Tao Chen, Li He, Pei-Yang Lin, Peng-Hu Wei, Guo-Guang Zhao, Yun-Zhe Liu

## Abstract

While invasive brain-computer interfaces have shown promise for high-performance speech decoding under medical use, the potential of intracranial stereoElectroEn-cephaloGraphy (sEEG), which causes less damage to patients, remains underexplored. With the rapid progress in representation learning, leveraging abundant pure recordings to further enhance speech decoding becomes increasingly attractive. However, some popular methods pre-train temporal models based on brain-level tokens, overlooking the brain’s desynchronization nature; others pre-train spatial-temporal models based on channel-level tokens, yet fail to evaluate them on more challenging tasks, e.g., speech decoding, which demands intricate processing in specific brain regions. To tackle these issues, we introduce a general pre-training framework for speech decoding – Du-IN-v2, which can extract contextual embeddings based on region-level tokens through discrete codex-guided mask modeling. To further push its limits, we propose Decoupling Product Quantization (DPQ), where different codexes are designed to extract different parts of brain dynamics. Our model achieves SOTA performance on both the 61-word classification task and the 49-syllable sequence classification task, surpassing all baselines. Model comparison and ablation studies reveal that our design choices, including (i) temporal modeling based on region-level tokens by utilizing 1D depthwise convolution to fuse channels in vSMC and STG regions and (ii) self-supervision by discrete decoupling codex-guided mask modeling, significantly contribute to these performances. Collectively, our approach, inspired by neuroscience findings, capitalizing on region-level representations from specific brain regions, is suitable for invasive brain modeling. It marks a promising neuro-inspired AI approach in BCI.

## 1 INTRODUCTION

Brain signals refer to the biometric information collected from the brain. Their patterns provide valuable insights toward understanding the physiological functions of the brain and the mechanism of related diseases, leading to various applications, including speech decoding (Cho et al., 2023; Duan et al., 2023; Moses et al., 2021), sleep cognition research (Liu & Jia, 2022; Zheng et al., 2023), neurological disorders detection (Jiang et al., 2024; Zhang et al., 2024), and so on. Due to the high signal-noise ratio, invasive recording methods (e.g., stereoElectroEncephaloGraphy (sEEG), ElectroCorticoGraphy (ECoG)) usually reveal these underlying mechanisms better than non-invasive recording methods. Many previous works (Jo et al., 2024; Duan et al., 2023) have shown that decoding speech from EEG signals is difficult, and the performance is limited. Compared with ECoG, sEEG imposes less trauma on patients and provides more stereotactic information from specific brain regions. Although some studies (Moses et al., 2021; Metzger et al., 2023) have recently shown promise for building high-performance speech decoders based on ECoG, there are few attempts made to explore the potential of sEEG-based speech decoding.

Modeling intracranial neural signals, especially sEEG, has drawn much research attention, but several issues remain unresolved. Current research on modeling neural signals is predominantly divided into two lines according to the basic modeling units (e.g., channel-level tokens or group-level tokens^1^). Some studies (Yuan et al., 2024; Jiang et al., 2024) utilize shared embedding blocks to embed single channels into channel-level tokens, neglecting the specificity of brain computation (Caucheteux et al., 2023); then they adopt spatial-temporal integration to model spatial relationships among them, attempting to regain the precise state of the brain. However, these methods primarily focus on channel-level classification tasks, e.g., seizure detection, yet fail to validate them on more challenging group-level classification tasks, e.g., speech decoding. Other studies (Duan et al., 2023; Feng et al., 2023) fuse all channels (across the whole brain) to build brain-level tokens, overlooking the brain’s desynchronization nature (Buzsaki, 2006); then they adopt temporal modeling to capture the rapid process of brain dynamics. Besides, labeling data at scale in medical experiments is often impractical or costly, emphasizing the need to maximize label efficiency. Therefore, developing an efficient pre-training framework that draws on prior neuroscience findings is highly appealing, as it can make the most of abundant unlabeled data.

The primary challenge in modeling intracranial neural signals lies in extracting meaningful tokens, requiring careful consideration of two key factors. (1) Temporal scale. Since intracranial neural signals have high temporal resolution and signal-noise ratio, these tokens must capture rapid dynamic changes in brain activity. (2) Spatial scale. Considering the brain’s desynchronization nature, these tokens should correctly capture the information of each brain region for further integration and, if needed, decouple different parts of brain dynamics within each brain region. To better assess how well different models capture the intricate processing within each brain region, we can evaluate these methods on tasks mainly involving a few brain regions.

Since speech mainly involves specific brain regions related to vocal production, as demonstrated in Section 2.1, we utilize speech decoding tasks to evaluate which model can effectively extract information from specific brain regions. Since there are too few open source sEEG language datasets (Angrick et al., 2021; Wang et al., 2023), we collected a well-annotated Chinese word-reading sEEG dataset (vocal production), including 12 subjects, which makes up for the problem of missing sEEG recordings in language tasks. Inspired by neuroscientific findings, we systematically demonstrate the locality and specificity of brain computation and propose the Du-IN-v2 model to solve the abovementioned issues. Compared to other existing methods for modeling brain signals, Du-IN-v2 achieves SOTA performance on both the 61-word classification task and the 49-syllable sequence classification task, demonstrating the effectiveness of our model in extracting meaningful tokens that can capture both the rapid changes and the precise state of specific brain regions. It marks a promising neuro-inspired AI approach (Saxe et al., 2021; Richards et al., 2019) in BCI.

To sum up, the main contributions of our work comprise:

1. A well-annotated Chinese word-reading sEEG dataset, addressing the lack of sEEG language dataset. The dataset will be publicly available.
2. Demonstration of brain-specific computation – achieving the best decoding performance only requires about one electrode in specific brain regions (i.e., vSMC, STG).
3. A novel framework for sEEG speech decoding – Du-IN-v2, which learns region-level contextual embeddings through discrete decoupling codex-guided mask modeling.
4. SOTA performance on the sEEG speech decoding task – Du-IN-v2 achieves top-1 accuracy of 64.53% on the 61-word classification task and 74.72% on the 49-syllable sequence classification task, surpassing all other baselines.

## 2 RELATED WORKS

### 2.1 NEURAL BASIS OF LANGUAGE FUNCTION

Neuroscientific research (Bouchard et al., 2013; Dichter et al., 2018; Sheng et al., 2019) in the past has extensively explored brain regions supporting language functionality. In neuroscience, the investigation into language functionality related to speech has been categorized into two main streams: one dedicated to semantic processing and the other to vocal production. Previous studies (Binder et al., 1997; Sheng et al., 2019) have shown that brain regions associated with semantic processing primarily include left inferior frontal gyrus (IFG), left anterior temporal lobe (ATL), and bilateral middle temporal gyrus (MTG).

As for vocal production, which is also the focus of our work, it is predominantly governed by motor information related to language articulation, primarily involving ventral sensorimotor cortex (vSMC), bilateral superior temporal gyrus (STG), and bilateral dorsal laryngeal motor cortex (dLMC) (Bouchard et al., 2013; Dichter et al., 2018; Chartier et al., 2018). Our analysis results based on our collected word-reading sEEG dataset also confirm this point, as illustrated in Figure 3.

### 2.2 LANGUAGE DECODING IN BCI

The keys to decoding natural language from brain signals are (1) high-quality recordings, and (2) well-designed models with good representations. Compared to non-invasive recordings (e.g., EEG), invasive recordings manifest advantages in providing detailed information about specific brain regions with a high signal-noise ratio. Since speech mainly involves some specific brain regions, obtaining detailed recordings of these brain regions will significantly enhance the decoding performance. Existing works (Cho et al., 2023; Moses et al., 2021; Feng et al., 2023) have shown the great potential of building a high-performance decoder based on invasive recordings.

The other key is well-designed models with good representations. Existing work for brain-to-language representations can be classified into two categories: self-supervision or alignment with representation models pre-trained on other modalities (e.g., text, audio). BrainBERT (Wang et al., 2023) learns general embeddings through self-supervised mask modeling. DeWave (Duan et al., 2023) introduces discrete codex encoding and aligns neural representations with text embeddings from BART (Lewis et al., 2019), thus enhancing the extraction of semantic processing-related information from EEG recordings. Metzger et al. (2023) align neural representations with acoustic embeddings to improve the extraction of vocal production-related information from ECoG recordings.

### 2.3 SELF-SUPERVISED LEARNING IN BCI

In recent years, self-supervised pre-training has made significant progress in natural language processing (Devlin et al., 2018; Radford et al., 2018; Brown et al., 2020) and computer vision (Bao et al., 2021; He et al., 2022; Chen et al., 2020). However, its potential in BCI is far from being explored. BrainBERT (for sEEG) (Wang et al., 2023) embeds single channels into channel-level tokens and utilizes mask modeling to learn general representations. Brant (for sEEG) (Zhang et al., 2024; Yuan et al., 2024), PopT (for sEEG) (Chau et al., 2024) and some works (for EEG) (Jiang et al., 2024; Guetschel et al., 2024) further adopt spatial-temporal integration to model spatial relationships among them. Some works (for EEG) (Kostas et al., 2021; Eldele et al., 2021; Wu et al., 2024; Foumani et al., 2024) take the other way – fusing all channels (across the whole brain) to build brain-level tokens, and it uses self-supervised learning to learn contextual representations. Considering the difference among brain regions, MMM (for EEG) (Yi et al., 2024) further splits channels into different groups to build region-level tokens.

All existing pre-training methods for sEEG primarily pre-train spatial-temporal models based on channel-level tokens yet only evaluate them on channel-level classification tasks, e.g., seizure detection. However, unlike EEG pre-training methods, their effectiveness over more challenging group-level classification tasks, e.g., speech decoding. Besides, there is no standard channel configuration for sEEG recordings, unlike EEG recordings, which makes modeling spatial relationships in sEEG more challenging.

## 3 METHOD

This paper aims to develop a general pre-training framework for speech decoding. The core idea is to use a VQ codex (Van Den Oord et al., 2017), enhanced by decoupling product quantization (Jegou et al., 2010), as a learnable neural vocabulary that can identify the precise states of specific brain regions. Once this vocabulary is optimized, we apply mask modeling to learn region-level contextual embeddings, enhancing performance on downstream tasks. The three-stage training framework of Du-IN-v2 is illustrated in Figure 1.

**Figure 1.**
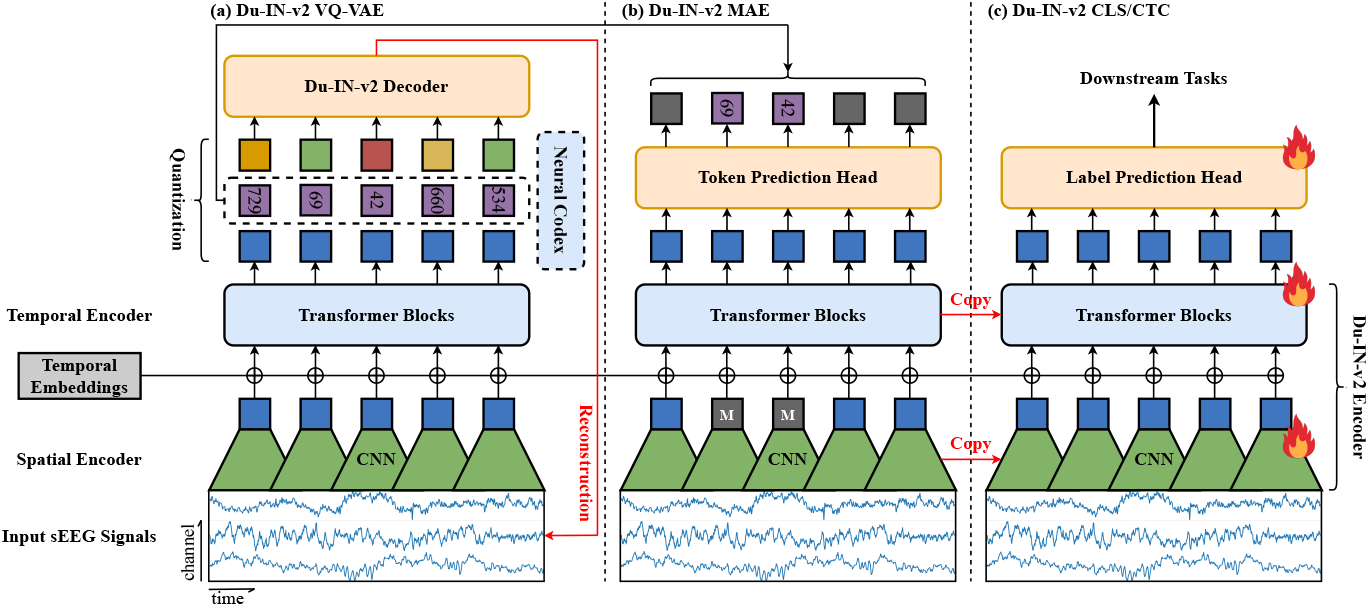
An overview of the three-stage training pipeline of Du-IN-v2. **(a)**.Learning discrete neural codex in the Du-IN-v2 VQ-VAE by reconstructing the original sEEG signals. **(b)**. Mask modeling pre-training of Du-IN-v2 Encoder in the Du-IN-v2 MAE. **(c)**. Fine-tuning the pre-trained Du-IN-v2 Encoder with an MLP head for various downstream speech decoding tasks.

### 3.1 TASK DEFINITION

Due to the lack of open-source sEEG datasets related to language tasks, we follow the experimental design outlined by Moses et al. (2021) to collect a well-annotated Chinese word-reading sEEG dataset (vocal production). During the experiment, each subject speaks aloud 61 pre-determined Chinese words 50 times; see Zheng et al. (2024) for more details. We formulate the multi-channel sEEG signals as 𝒳∈ ℝ^*C×T*^, where *C* is the number of sEEG channels and *T* is the total timestamps. The associated label is denoted as ***y*** ∈ 𝒴, where 𝒴 represents either the set of 61 pre-determined words or the corresponding syllable sequences, depending on the downstream task; see Appendix B for more details. In summary, this dataset comprises paired sEEG-label data (⟨𝒳, *y* ⟩), and we develop different Du-IN-v2 variants to decode the corresponding label ***y*** from a sequence of raw sEEG signals 𝒳– Du-IN-v2 CLS for the 61-word classification task, Du-IN-v2 CTC for the 49-syllable sequence classification task.

### 3.2 MODEL ARCHITECTURE

We introduce the Du-IN-v2 Encoder, a general architecture for sEEG speech decoding tasks, which is commonly used in all Du-IN-v2 variants for feature extraction, as shown in Figure 1. The Du-IN-v2 Encoder consists of two parts: (1) Spatial Encoder and (2) Temporal Encoder.

#### Spatial Encoder

As each sEEG sample 𝒳 has multiple channels, it is vital to fuse different channels to extract meaningful tokens before token-wise interaction by self-attention. We employ a spatial encoder, which consists of a linear projection and several convolution blocks, to encode each sEEG sample into tokens. The linear projection transforms the raw sEEG signals into the hidden neural space, and its weights are utilized for subsequent analysis. The convolution block is composed of a 1-D depthwise convolution layer, a group normalization layer (Wu & He, 2018), and a GELU activation function (Hendrycks & Gimpel, 2016). To avoid the loss of boundary information (Huang et al., 2024), we apply convolution blocks on patches with overlap to generate the token sequence, which can better capture the rapid dynamics changes in brain activity. We denote the output tokens from the spatial encoder as

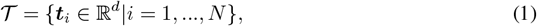

where *D* is the dimension of tokens (and embeddings) and *N* is the number of tokens.

#### Temporal Embedding

In order to enable the model to be aware of the temporal information of tokens, we utilize the parameter-free position embeddings introduced by Vaswani et al. (2017), i.e., 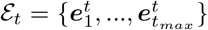 Note that *t*_*max*_ is the hyperparameter determining the maximum number of time steps and *t*_*max*_ ≥ *N*. Given one arbitrary token ***t***_*i*_ in Equation 1 from the spatial encoder, we add the corresponding temporal embedding to it:

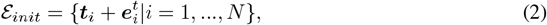

which forms the input embeddings E_*init*_ for the Transformer Encoder.

#### Temporal Encoder

Finally, the sequence of embeddings will be directly fed into the Transformer encoder (Vaswani et al., 2017) to get the final encoded ε = {***e***_*i*_ ∈ ℝ^*d*^| *i* = 1, …, *N*}. To make the training of the Transformer more stable and efficient, we incorporate some modifications introduced by Dehghani et al. (2023). We add layer normalization to the queries and keys before the dot-product attention mechanism, which avoids over-large values in attention logits:

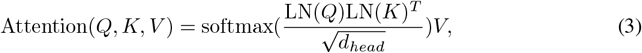

where *d*_*head*_ is the dimension of attention head and LN denotes layer normalization (Ba et al., 2016).

### 3.3 DU-IN-V2 VQ-VAE TRAINING

Prior to pre-training Du-IN-v2 through mask modeling (i.e., Du-IN-v2 MAE training), we need to discretize sEEG tokens into discrete codes. We introduce vector-quantized neural signal regression, which is trained by reconstructing the original sEEG signals, as shown in Figure 1 (a). The key components are (1) Du-IN-v2 Encoder, (2) Vector Quantizer, and (3) Du-IN-v2 Decoder. The idea is basically inspired by VQ-VAE (Van Den Oord et al., 2017), which encodes images into discrete codes. To better capture the micro-scale functional modules within certain brain regions (Chapeton et al., 2022), we introduce Decoupling Product Quantization (DPQ) (Jegou et al., 2010), as shown in Figure 2. In this approach, different codexes are designed to extract different parts of brain dynamics.

**Figure 2.**
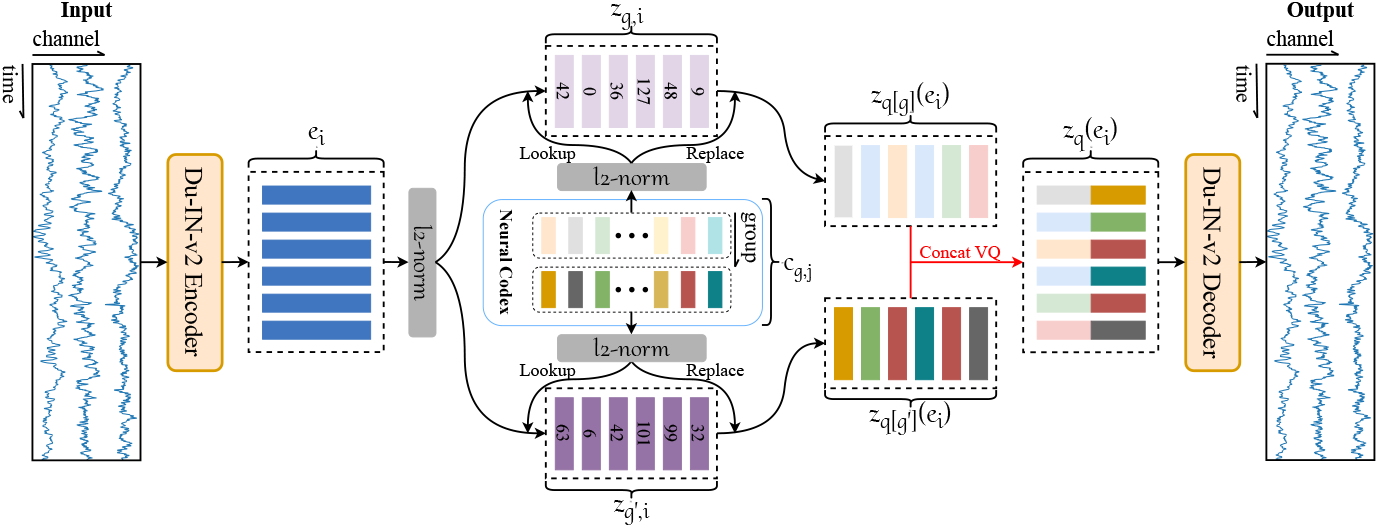
An illustration of Decoupling Product Quantization (DPQ) for Du-IN-v2 framework. In practice, we instantiate DPQ with 4 parallel neural codexes (each consisting of 128 discrete codes).

**Figure 3.**
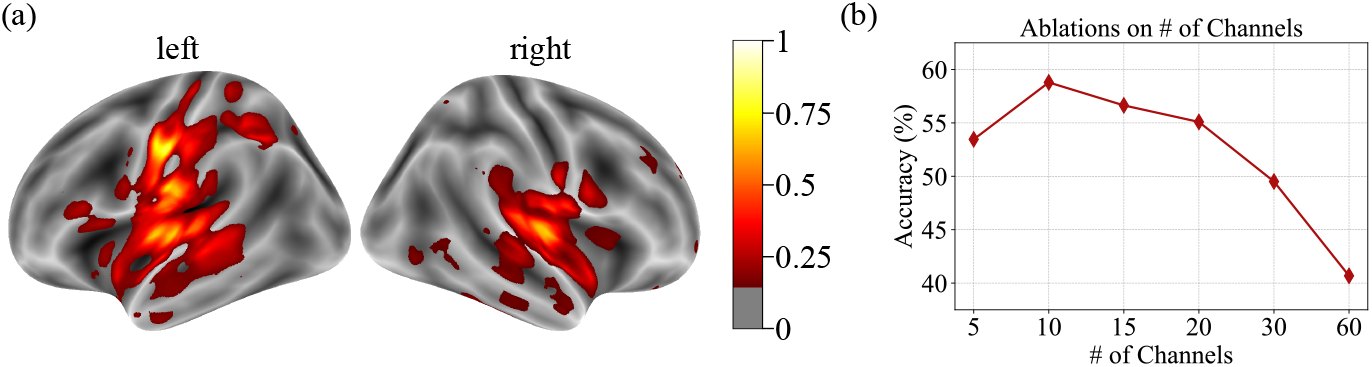
The channel contribution analysis. **(a)**.The channel contribution map. **(b)**. The effect of the number of channels (sorted by channel contribution scores) on word classification performance.

#### Vector Quantizer

The output embeddings ε = {*e*_*i*_ ∈ ℝ^*d*^| *i* = 1, …, *N*} from the Du-IN-v2 Encoder are fed into a vector quantizer, which consists of *G* parallel sub-quantizers (i.e., neural codexes). The *g*-th neural codex is defined as 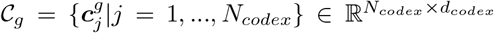, where *N*_*codex*_ is the number of the discrete neural codes and *d*_*codex*_ is the dimension of each code embedding. After that, we utilize a linear projection 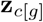 to get the mapped embeddings 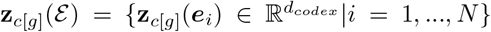 in the codex space. Then, the codex looks up the nearest neighbor of each embedding 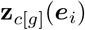 in the neural codex 𝒞_*g*_. This procedure can be formulated as

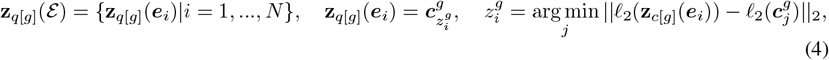

where *ℓ*_2_ represents *ℓ*_2_ normalization and 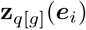 is the quantized vector after the *g*-th sub-quantizer. This is equivalent to finding the closest neural embedding by cosine similarity, and such *ℓ*_2_ normalization improves the codex utilization (Peng et al., 2022). As illustrated in Figure 2, the output 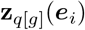 of the *G* sub-quantizers is concatenated to the full code 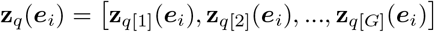. Then, the code z_*q*_(*e*_*i*_) is linearly mapped to the vector-quantized embedding *z*_*i*_ ∈ ℝ^*d*^.

#### Du-IN-v2 Decoder

The Du-IN-v2 Decoder consists of a Transformer decoder and a stack of 1D transposed convolution layers. Given a sequence of the vector-quantized embeddings *Ƶ* = {***z***_*i*_|*i* = 1, …, *N*}, the Du-IN-v2 Decoder convert these discrete embeddings back into raw sEEG signals 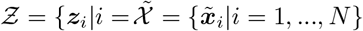. The mean squared error (MSE) loss is utilized to guide the regression. The total loss for training the Du-IN-v2 VQ-VAE model is defined as:

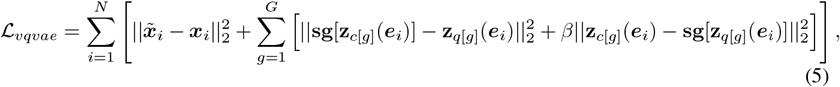

where **sg** represents the stop-gradient operation, which is an identity at the forward pass and has zero gradients. To stabilize the codex update, we use the exponential moving average strategy (Van Den Oord et al., 2017).

### 3.4 DU-IN-V2 MAE TRAINING

#### Masked sEEG Modeling

To enforce Du-IN-v2 learning contextual representations, we propose masked sEEG modeling. The whole procedure is presented in Figure 1 (b). Given a sEEG sample 𝒳, the spatial encoder first transforms it to tokens 𝒯 = {*t*_*i*_|*i* = 1, …, *N*}. Given these tokens 𝒯, we adopt the consecutive mask strategy introduced by Baevski et al. (2020) with mask span as 3, and around 50% of tokens are masked. The masked position is termed as ℳ. Then, a shared learnable embedding *e*_[*M*]_ ∈ ℝ^*d*^ is used to replace the original tokens:

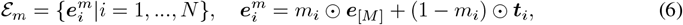

where *δ*() is the indicator function and *m*_*i*_ = *δ*(*I* ∈ *ℳ*). After that, the masked embeddings ε_*m*_ will be added by temporal embeddings ε_*t*_, and then fed into the Transformer encoder. The output embeddings ε will be used to predict the indices of the corresponding codes from the codex in the Du-IN-v2 VQ-VAE through a linear classifier:

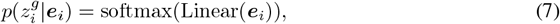

The training loss of mask modeling is defined as:

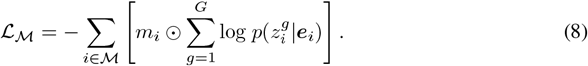

#### Symmetric Masking

Inspired by LaBraM (Jiang et al., 2024), we further introduce a symmetric masking strategy to improve training efficiency. We calculate the inverse of the generated mask ℳ, obtaining 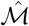. Similarly, we use the new mask 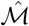 to perform the mask modeling, obtaining the mask modeling loss 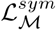. The total loss for training the Du-IN-v2 MAE model is defined as:

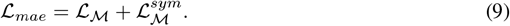

### 3.5 DU-IN-V2 CLS/CTC FINE-TUNING

We develop different Du-IN-v2 variants for different downstream tasks – the Du-IN-v2 CLS model for the 61-word classification task and the Du-IN-v2 CTC model for the 49-syllable sequence classification task. The Du-IN-v2 Encoder in these Du-IN-v2 variants is loaded from the pre-trained Du-IN-v2 MAE model. All parameters in Du-IN-v2 variants are trainable during fine-tuning.

#### Du-IN-v2 CLS model

The Du-IN-v2 CLS model adds a randomly initialized MLP on top of the Du-IN-v2 Encoder to decode the embeddings ε into the word label from 61 pre-determined words; see Appendix B.3 for more details. Following Moses et al. (2021), the Du-IN-v2 CLS model is optimized by minimizing a cross-entropy loss.

#### Du-IN-v2 CTC model

The Du-IN-v2 CTC model adds a randomly initialized MLP on top of the Du-IN-v2 Encoderto decode the embeddings ε into the syllable label sequence from 49 pre-defined syllables; see Appendix B.4 for more details. Following Metzger et al. (2023), the Du-IN-v2 CTC model is optimized by minimizing a CTC loss (Graves et al., 2006).

## 4 EXPERIMENTS

### 4.1 DATASET

Due to the lack of open-source sEEG datasets related to language tasks, we follow the experimental design outlined by Moses et al. (2021) to collect a well-annotated Chinese word-reading sEEG dataset (vocal production), including 12 subjects. The subjects undergo a surgical procedure to implant 7 to 13 invasive sEEG electrodes, each with 72 to 158 channels, in their brain. For each subject, the dataset contains 15 hours of 2000Hz recordings, 3 hours of which are task recordings.

#### Pre-training dataset

For each subject, the pre-training dataset contains all sEEG recordings (with about 54 million timestamps) of that subject. To stabilize computing resource usage, the time length of sEEG sample X is set to 4 seconds.

#### Downstream dataset

For each subject, 3 hours of the sEEG recordings are task recordings. The sEEG signals are segmented into about 3000 3-second samples, each of which is paired with the corresponding label (from either 61 pre-determined words or the corresponding syllable sequences).

### 4.2 IMPLEMENTATION DETAILS

#### Preprocess

We first filter the sEEG signals between 0.5Hz and 200Hz to remove low-frequency noise. Then, a notch filter of 50Hz is applied to avoid power-line interference. After that, all sEEG signals are resampled to 400Hz and bi-polar re-referenced (Li et al., 2018). Finally, we perform z-score normalization on each channel to guarantee normalized data scales across all channels.

#### Model Configurations

The “Spatial Encoder” contains one linear projection and three 1-D depthwise convolution layers, transforming the raw sEEG signals into tokens with *d* = 256. As the convolution along the time axis reduces temporal resolution, this process generates 40 tokens per sample in the pre-training dataset and 30 tokens per sample in the downstream dataset. The following “Transformer Encoder” contains an 8-layer Transformer encoder with model dimension *d* = 256, inner dimension (FFN) *d*_*ff*_ = 1024, and 8 attention heads. See Appendix B for more details.

#### Pre-training

During the pre-training, we use either all sEEG recordings (15 hours) or the sEEG recordings without task recordings (12 hours) to train the Du-IN-v2 VQ-VAE and Du-IN-v2 MAE models. To enhance the robustness of the learned codex and representations, we further use data augmentation described in Appendix C. For each subject, the model is pre-trained on a Linux system with 2 CPUs (Intel Xeon Gold 6230 40-Core Processor) and 1 GPU (NVIDIA Tesla V100 32GB) for ∼ 1.2 days.

#### Fine-tuning

During the downstream evaluation, we split the task recordings into training, validation, and testing splits with a size roughly proportional to 80%, 10%, and 10%. All experiments are conducted on the same machine with the same set of random seeds. The train/validation/test splits are the same across different models. We also use data augmentation, as described in Appendix C, to make the most of the gathered dataset. We employ cross-entropy loss (multi-class classification) as the training loss. Our experiments are conducted on one V100 GPU by Python 3.11.7 and PyTorch 2.1.2 + CUDA 12.3. The best models are trained based on the training set, selected from the validation set according to accuracy, and finally evaluated on the test set. For model comparison, we report the average and standard error values (of all subjects) on six different random seeds to obtain comparable results. For the results of the subject-wise evaluation, we report the average and standard deviation values (of each subject) in Appendix I.

### 4.3 CHANNEL CONTRIBUTION AND SELECTION

As demonstrated in Section 2.1, previous neuroscience studies reveal that vocal production predominantly engages specific brain regions. Given the sparse distribution of implanted sEEG electrodes (each containing 8-16 channels), it’s vital to exclude redundant electrodes unrelated to vocal production, thus improving decoding performance. We retain electrodes implanted in relevant brain regions and evaluate the performance based on the remaining electrodes. Table 1 demonstrates that excluding approximately 85% electrodes even leads to a dramatic increase in decoding performance.

**Table 1:** The word classification performance of Du-IN-v2 with or without electrode selection.

| Methods | # of Channels (Averaged) | Accuracy (%) ± ste (%) |
| --- | --- | --- |
| **Du-IN-v2** (w/o electrode selection) | 109.75 | 32.44±4.84 |
| **Du-IN-v2** (w/ electrode selection) | 12.25 | 58.02±4.02 |

To further understand the detailed contribution of each channel, we analyze the weights of linear projection in the spatial encoder. In detail, we calculate the contribution scores of channels per subject and organize them accordingly, as described in Appendix G. Figure 3 demonstrates that (1) the brain regions effective for speech decoding align with findings from previous neuroscience research, and (2) our model achieves optimal decoding performance with approximately 10 channels, 80% of which originate from the same electrode. To streamline, we utilize these top 10 channels (selected according to train-set) for both pre-training and downstream evaluation.

### 4.4 COMPARASION WITH OTHER MODELS

Table 2 presents the results of our Du-IN-v2 model and the advanced baselines that are designed for either brain signals or general time series. See Appendix A and Appendix B for detailed descriptions of models. The results demonstrate that our Du-IN-v2 model outperforms all baselines. It’s worth noting that the models (i.e., the foundation models designed for brain signals) that adopt spatialtemporal integration to model spatial relationships among channel-level tokens perform worse than the models that adopt temporal modeling based on region-level tokens, challenging the generalizability of current strategies to model spatial relationships among channels with Transformer.

**Table 2:**
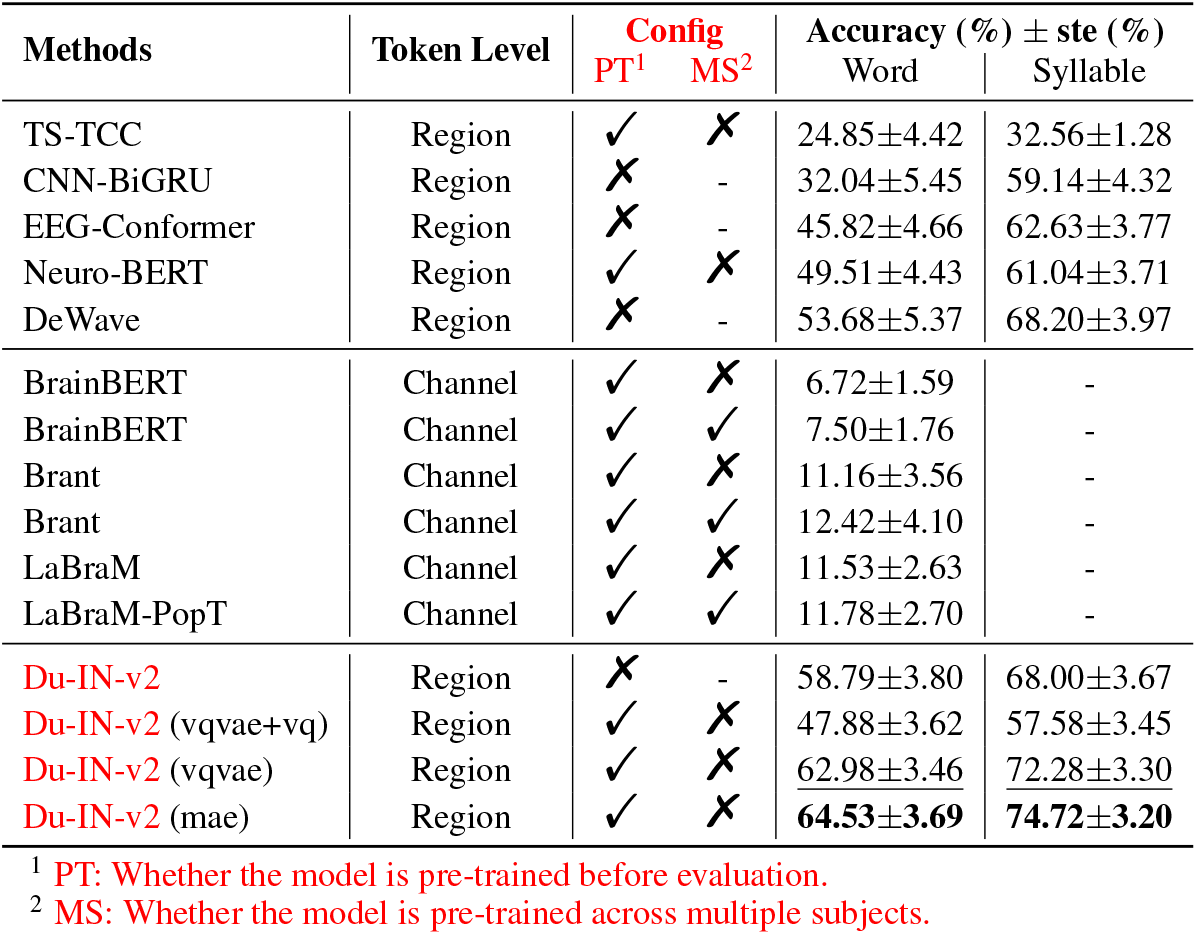
The performance of different methods (with the best in bold and the second underlined).

As BrainBERT (Wang et al., 2023) doesn’t consider the spatial relationships among channels, we mainly focus on understanding why Brant (Zhang et al., 2024), LaBraM (Jiang et al., 2024) and LaBraM-Popt (Jiang et al., 2024; Chau et al., 2024) fail to effectively capture the discriminative features on the speech decoding task. These models typically build channel-level tokens by segmenting non-overlapping patches with large receptive fields (e.g., 1 second) from single channels. However, this approach makes it challenging to capture the rapid process of brain dynamics. Since these models operate at a coarse time scale, they cannot be adapted to handle the 49-syllable sequence classification task. Moreover, while these models further utilize Transformer to capture the spatial relationships among these tokens, they do not encourage region-level embeddings, either through their architecture (Yi et al., 2024) or their pre-training objective (Chau et al., 2024). Therefore, the effectiveness of building brain foundation models based on these spatial-temporal backbones is still under exploration, especially for cognitive tasks (e.g., speech decoding), which are of great value in the field of neuroscience.

For other baselines that use temporal modeling based on region-level tokens, we provide a detailed explanation of their performance differences as follows. TS-TCC (Eldele et al., 2021) tokenizes raw sEEG signals into region-level tokens with a stack of 1D depthwise convolution blocks, but it lacks a temporal Transformer for further integration over time. CNN-BiGRU (Moses et al., 2021) introduces a stack of GRU layers on top of these tokens to perform temporal integration. Neuro-BERT (Wu et al., 2024) introduces a temporal Transformer to better integrate global temporal information. Compared to our Du-IN-v2 model, Neuro-BERT performs worse because length-fixed patches are prone to losing temporal boundary information (Huang et al., 2024). EEG-Conformer (Song et al., 2022) tokenizes raw sEEG signals with the temporal-spatial convolution, applying the same convolutional kernel across different channels, which overlooks the specificity of brain computation (Caucheteux et al., 2023). This also raises a challenge for the effectiveness of current sEEG foundation models, which rely on shared convolution blocks across individual channels. DeWave (Duan et al., 2023) utilizes the Conformer model (Gulati et al., 2020) for tokenization, which involves more parameters but is less effective than 1D depthwise convolution.

### 4.5 ABLATION STUDY

#### Self-Supervision Initialization

As illustrated in Figure 1, the Du-IN-v2 model entails a two-stage pre-training process, wherein both the Du-IN-v2 VQ-VAE model and the Du-IN-v2 MAE model are trained. Previous studies utilize different strategies (Duan et al., 2023; Chen et al., 2024; Jiang et al., 2024) to leverage these pre-trained models to enhance the performance of downstream tasks. Here, we evaluate these different strategies for comparison; see Appendix B.3 for detailed definitions. Table 2 shows that initializing weights from the Du-IN-v2 MAE model captures contextual embeddings effectively, resulting in the highest decoding performance.

#### Pre-training with/without Downstream Datasets

During the pre-training stage, we hope that the Du-IN-v2 VQ-VAE model can extract general tokens of that brain region, thus guiding the Du-IN-v2 MAE model to learn general representations that are not specific to any particular task. Although no label data is used during the pre-training stage, to eliminate the influence of the pre-training data on downstream tasks, we compare the results with or without incorporating the downstream task dataset into the pre-training stage. Table 3 shows a slight performance drop when excluding downstream datasets. However, the decoding performance is still higher than the baseline performance without pre-training, which means that the degradation is mainly due to the decrease of the pre-training dataset. We hope that, with more pure recordings, our model can achieve better decoding performance.

**Table 3:** Ablation study on whether pre-training with the downstream dataset (DD) or not.

| Methods | Pre-training Dataset Size | Accuracy (%) ± ste (%) | |
| --- | --- | --- | --- |
|  |  | word | syllable |
| Du-IN-v2 (mae w/o DD) | 12 hours per subject | 62.19±3.74 | 71.94±3.36 |
| Du-IN-v2 (mae w/ DD) | 15 hours per subject | 64.53±3.69 | 74.72±3.20 |

**Table 4:**
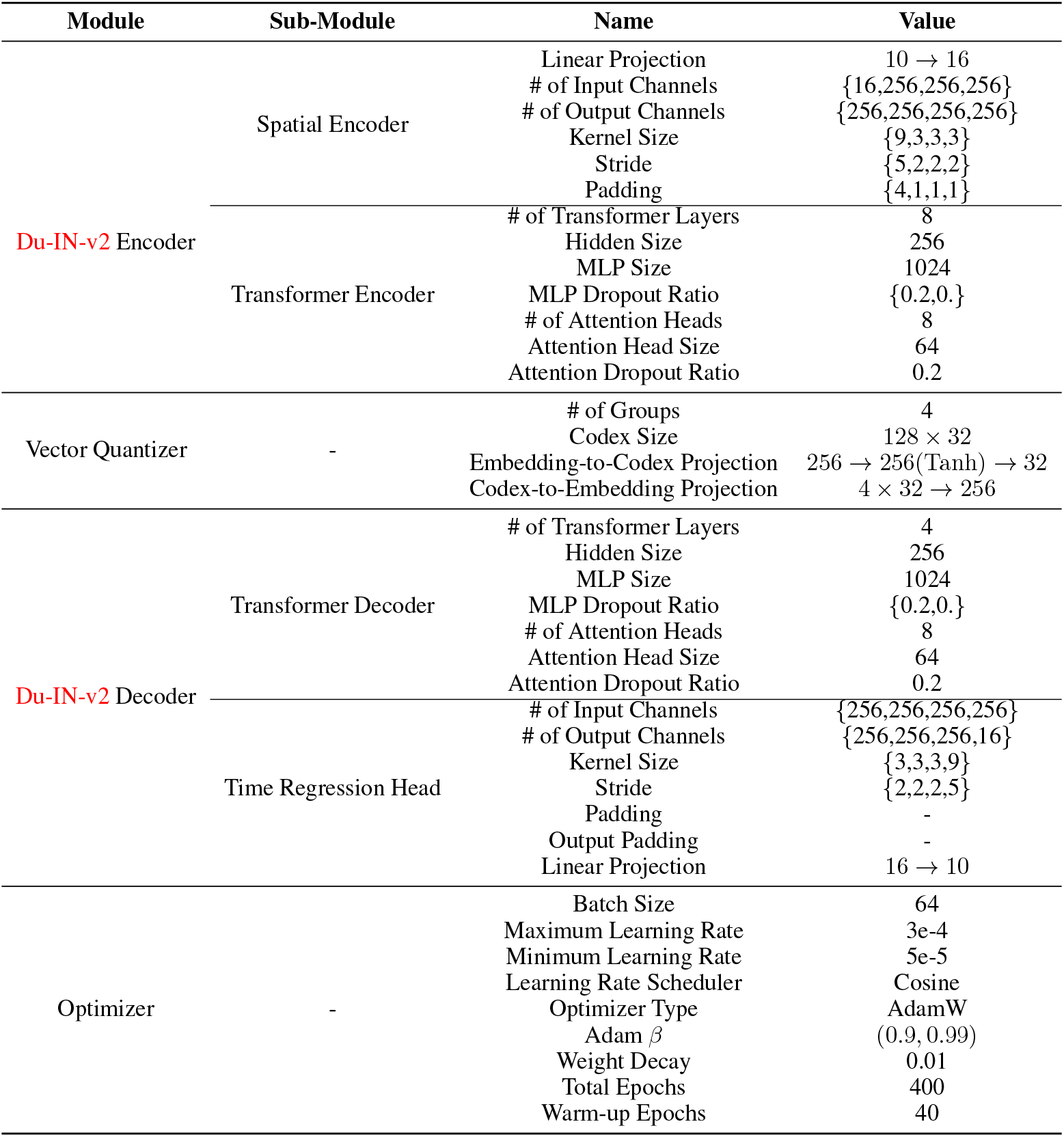
The hyperparameters for Du-IN-v2 VQ-VAE training.

**Table 5:** The hyperparameters for Du-IN-v2 MAE training.

| Module | Sub-Module | Name | Value |
| --- | --- | --- | --- |
| Token Prediction Head | - | # of Prediction Heads | 4 |
| | | Linear Projection | 256 → 128 |
| Optimizer | - | Batch Size | 64 |
|  |  | Maximum Learning Rate | 3e-4 |
|  |  | Minimum Learning Rate | 5e-5 |
|  |  | Learning Rate Scheduler | Cosine |
|  |  | Optimizer Type | AdamW |
| | | Adam $\beta$ | (0.9, 0.99) |
|  |  | Weight Decay | 0.05 |
|  |  | Total Epochs | 400 |
|  |  | Warm-up Epochs | 40 |

**Table 6:**
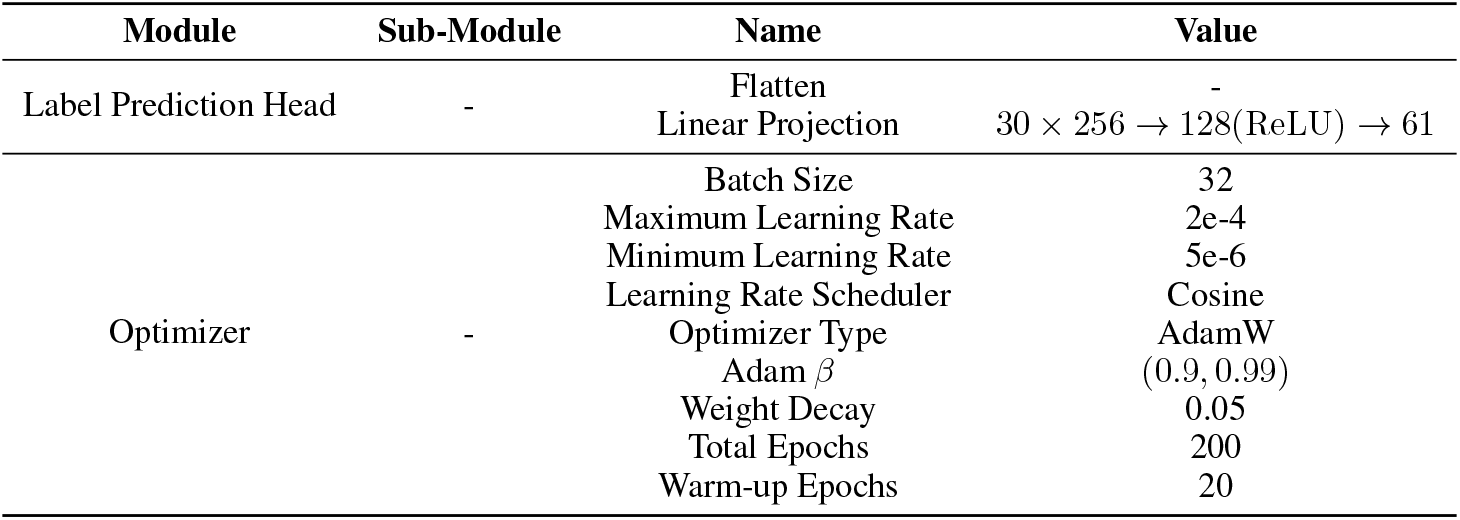
The hyperparameters for Du-IN-v2 CLS training.

**Table 7:** The hyperparameters for Du-IN-v2 CTC training.

| Module | Sub-Module | Name | Value |
| --- | --- | --- | --- |
| Label Prediction Head | - | Flatten Window | 3 |
| | | Linear Projection | $3 \times 256 \rightarrow 128(\text{ReLU}) \rightarrow 49$ |
| Optimizer | - | Batch Size | 32 |
|  |  | Maximum Learning Rate | 2e-4 |
|  |  | Minimum Learning Rate | 5e-6 |
|  |  | Learning Rate Scheduler | Cosine |
|  |  | Optimizer Type | AdamW |
| | | Adam $\beta$ | (0.9, 0.99) |
|  |  | Weight Decay | 0.05 |
|  |  | Total Epochs | 200 |
|  |  | Warm-up Epochs | 20 |

**Table 8:** Ablations to validate the effectiveness of region-specific channel selection.

| Settings | Setting 1 | Setting 2 | Setting 3 |
| --- | --- | --- | --- |
| MSE | **0.2969±0.0376** | 0.5211±0.0492 | 0.9673±0.0148 |

<sup>1</sup> Setting 1: Select the top 10 speech-related channels for neural signal reconstruction.
<sup>2</sup> Setting 2: Randomly select 10 channels for neural signal reconstruction.
<sup>3</sup> Setting 3: Use all channels (109.75 channels on average) for neural signal reconstruction.

**Table 9:**
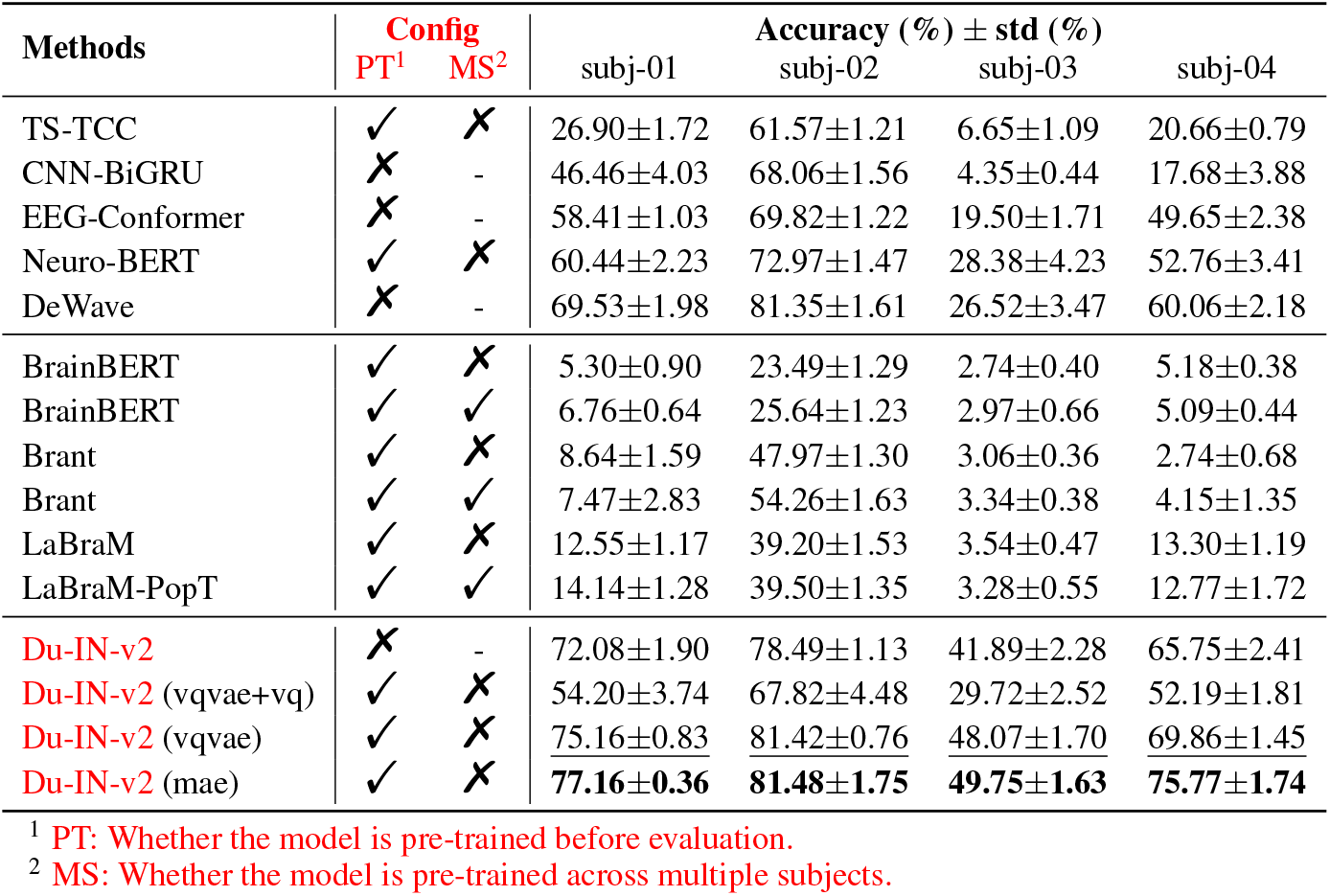
The 61-word performance of different methods from subjects (01-04).

**Table 10:** The 61-word performance of different methods from subjects (05-08).

| Methods | Config | | Accuracy (%) ± std (%) | | | |
| --- | --- | --- | --- | --- | --- | --- |
|  | PT<sup>1</sup> | MS<sup>2</sup> | subj-05 | subj-06 | subj-07 | subj-08 |
| TS-TCC | ✓ | ✗ | 34.53±1.34 | 9.73±0.97 | 24.83±1.15 | 20.08±1.60 |
| CNN-BiGRU | ✗ | - | 51.26±4.93 | 31.52±1.48 | 47.75±1.12 | 24.64±4.44 |
| EEG-Conformer | ✗ | - | 65.44±1.31 | 31.06±2.58 | 47.89±1.86 | 42.12±2.08 |
| Neuro-BERT | ✓ | ✗ | 71.61±1.97 | 36.63±2.15 | 50.23±2.63 | 43.24±1.36 |
| DeWave | ✗ | - | 73.44±0.66 | 35.17±1.93 | 51.93±1.18 | 52.89±1.44 |
| BrainBERT | ✓ | ✗ | 9.59±0.90 | 2.67±0.31 | 4.79±0.64 | 5.10±0.59 |
| BrainBERT | ✓ | ✓ | 11.28±1.17 | 3.00±0.47 | 5.31±0.55 | 5.22±0.76 |
| Brant | ✓ | ✗ | 24.10±2.14 | 5.09±1.27 | 7.74±1.46 | 8.66±1.07 |
| Brant | ✓ | ✓ | 28.83±1.93 | 5.28±1.48 | 9.70±1.85 | 8.93±1.39 |
| LaBraM | ✓ | ✗ | 15.52±1.48 | 6.63±0.33 | 12.65±0.50 | 7.41±0.86 |
| LaBraM-PopT | ✓ | ✓ | 17.73±1.11 | 5.81±1.61 | 12.90±1.26 | 6.41±1.85 |
| Du-IN-v2 | ✗ | - | 70.52±1.18 | 43.30±1.52 | 58.69±0.88 | 55.10±1.27 |
| Du-IN-v2 (vqvae+vq) | ✓ | ✗ | 62.79±2.24 | 35.29±1.62 | 47.91±2.74 | 44.02±1.34 |
| Du-IN-v2 (vqvae) | ✓ | ✗ | 73.28±1.45 | <u>46.94±0.52</u> | <u>61.10±2.90</u> | <u>58.42±1.37</u> |
| Du-IN-v2 (mae) | ✓ | ✗ | **74.12±0.55** | **49.71±1.02** | **66.78±1.80** | **59.04±1.16** |

<sup>1</sup> PT: Whether the model is pre-trained before evaluation.<sup>2</sup> MS: Whether the model is pre-trained across multiple subjects.

**Table 11:** The 61-word performance of different methods from subjects (09-12).

| Methods | Config | | Accuracy (%) ± std (%) | | | |
| --- | --- | --- | --- | --- | --- | --- |
|  | PT<sup>1</sup> | MS<sup>2</sup> | subj-09 | subj-10 | subj-11 | subj-12 |
| TS-TCC | ✓ | ✗ | 37.75±1.22 | 5.71±0.38 | 35.72±0.95 | 14.12±0.68 |
| CNN-BiGRU | ✗ | - | 44.03±5.88 | 7.11±0.71 | 28.44±3.42 | 13.17±3.41 |
| EEG-Conformer | ✗ | - | 56.51±1.98 | 22.22±1.07 | 57.10±2.03 | 29.87±1.44 |
| Neuro-BERT | ✓ | ✗ | 54.12±4.11 | 24.66±1.28 | 62.99±0.93 | 36.07±2.36 |
| DeWave | ✗ | - | 61.18±1.10 | 19.75±1.20 | 70.25±1.97 | 42.08±6.05 |
| BrainBERT | ✓ | ✗ | 6.82±1.42 | 2.55±0.49 | 8.73±0.92 | 3.71±1.02 |
| BrainBERT | ✓ | ✓ | 7.20±1.37 | 2.49±0.43 | 10.60±1.22 | 4.41±0.88 |
| Brant | ✓ | ✗ | 6.82±1.44 | 2.84±0.26 | 8.76±1.55 | 7.53±1.29 |
| Brant | ✓ | ✓ | 6.46±1.66 | 3.00±0.31 | 9.82±1.71 | 7.82±1.66 |
| LaBraM | ✓ | ✗ | 8.97±0.52 | 3.50±0.30 | 7.92±0.61 | 7.19±0.54 |
| LaBraM-PopT | ✓ | ✓ | 9.35±1.09 | 3.91±0.31 | 7.84±0.92 | 7.74±1.24 |
| Du-IN-v2 | ✗ | - | 60.14±2.82 | 32.62±1.38 | 68.75±2.40 | 58.20±2.98 |
| Du-IN-v2 (vqvae+vq) | ✓ | ✗ | 56.71±1.48 | 24.07±1.59 | 54.82±3.78 | 45.07±2.56 |
| Du-IN-v2 (vqvae) | ✓ | ✗ | 65.79±1.18 | **41.30±1.45** | <u>72.86±1.45</u> | **61.61±1.95** |
| Du-IN-v2 (mae) | ✓ | ✗ | **66.83±1.42** | <u>37.85±1.01</u> | **74.38±1.40** | <u>61.43±1.42</u> |

<sup>1</sup> PT: Whether the model is pre-trained before evaluation.<sup>2</sup> MS: Whether the model is pre-trained across multiple subjects.

**Table 12:**
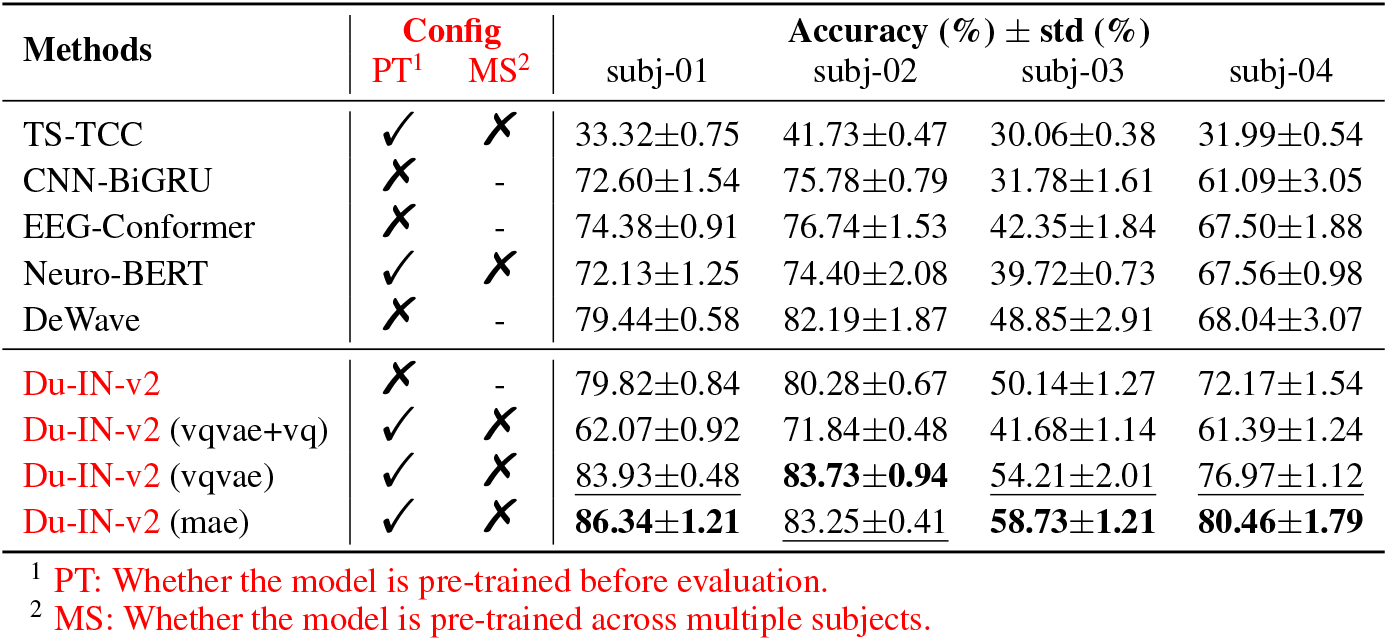
The 49-syllable performance of different methods from subjects (01-04).

| Methods | Config | | Accuracy (%) ± std (%) | | | |
| --- | --- | --- | --- | --- | --- | --- |
|  | PT<sup>1</sup> | MS<sup>2</sup> | subj-01 | subj-02 | subj-03 | subj-04 |
| TS-TCC | ✓ | ✗ | 33.32±0.75 | 41.73±0.47 | 30.06±0.38 | 31.99±0.54 |
| CNN-BiGRU | ✗ | - | 72.60±1.54 | 75.78±0.79 | 31.78±1.61 | 61.09±3.05 |
| EEG-Conformer | ✗ | - | 74.38±0.91 | 76.74±1.53 | 42.35±1.84 | 67.50±1.88 |
| Neuro-BERT | ✓ | ✗ | 72.13±1.25 | 74.40±2.08 | 39.72±0.73 | 67.56±0.98 |
| DeWave | ✗ | - | 79.44±0.58 | 82.19±1.87 | 48.85±2.91 | 68.04±3.07 |
| Du-IN-v2 | ✗ | - | 79.82±0.84 | 80.28±0.67 | 50.14±1.27 | 72.17±1.54 |
| Du-IN-v2 (vqvae+vq) | ✓ | ✗ | 62.07±0.92 | 71.84±0.48 | 41.68±1.14 | 61.39±1.24 |
| Du-IN-v2 (vqvae) | ✓ | ✗ | 83.93±0.48 | **83.73±0.94** | <u>54.21±2.01</u> | <u>76.97±1.12</u> |
| Du-IN-v2 (mae) | ✓ | ✗ | **86.34±1.21** | <u>83.25±0.41</u> | **58.73±1.21** | **80.46±1.79** |

<sup>1</sup> PT: Whether the model is pre-trained before evaluation.
<sup>2</sup> MS: Whether the model is pre-trained across multiple subjects.

**Table 13:** The 49-syllable performance of different methods from subjects (05-08).

| Methods | Config | | Accuracy (%) ± std (%) | | | |
| --- | --- | --- | --- | --- | --- | --- |
|  | PT<sup>1</sup> | MS<sup>2</sup> | subj-05 | subj-06 | subj-07 | subj-08 |
| TS-TCC | ✓ | ✗ | 36.73±1.17 | 29.50±0.54 | 30.06±0.78 | 32.85±0.91 |
| CNN-BiGRU | ✗ | - | 80.31±1.73 | 51.19±1.46 | 52.81±1.26 | 55.79±1.63 |
| EEG-Conformer | ✗ | - | 80.43±0.77 | 50.34±1.58 | 57.13±2.27 | 58.97±1.13 |
| Neuro-BERT | ✓ | ✗ | 78.23±2.06 | 51.64±1.08 | 52.25±2.25 | 54.20±1.92 |
| DeWave | ✗ | - | 85.95±1.52 | 59.88±0.36 | 61.29±2.57 | 66.41±0.81 |
| Du-IN-v2 | ✗ | - | 84.53±1.07 | 57.51±1.40 | 62.01±1.71 | 63.97±1.82 |
| Du-IN-v2 (vqvae+vq) | ✓ | ✗ | 75.16±1.15 | 48.72±1.34 | 54.38±0.85 | 51.97±1.59 |
| Du-IN-v2 (vqvae) | ✓ | ✗ | <u>86.64±1.44</u> | <u>64.74±0.55</u> | <u>66.61±1.07</u> | <u>67.36±0.68</u> |
| Du-IN-v2 (mae) | ✓ | ✗ | **87.66±1.26** | **66.20±1.12** | **71.53±0.27** | **70.65±1.32** |

<sup>1</sup> PT: Whether the model is pre-trained before evaluation.
<sup>2</sup> MS: Whether the model is pre-trained across multiple subjects.

**Table 14:**
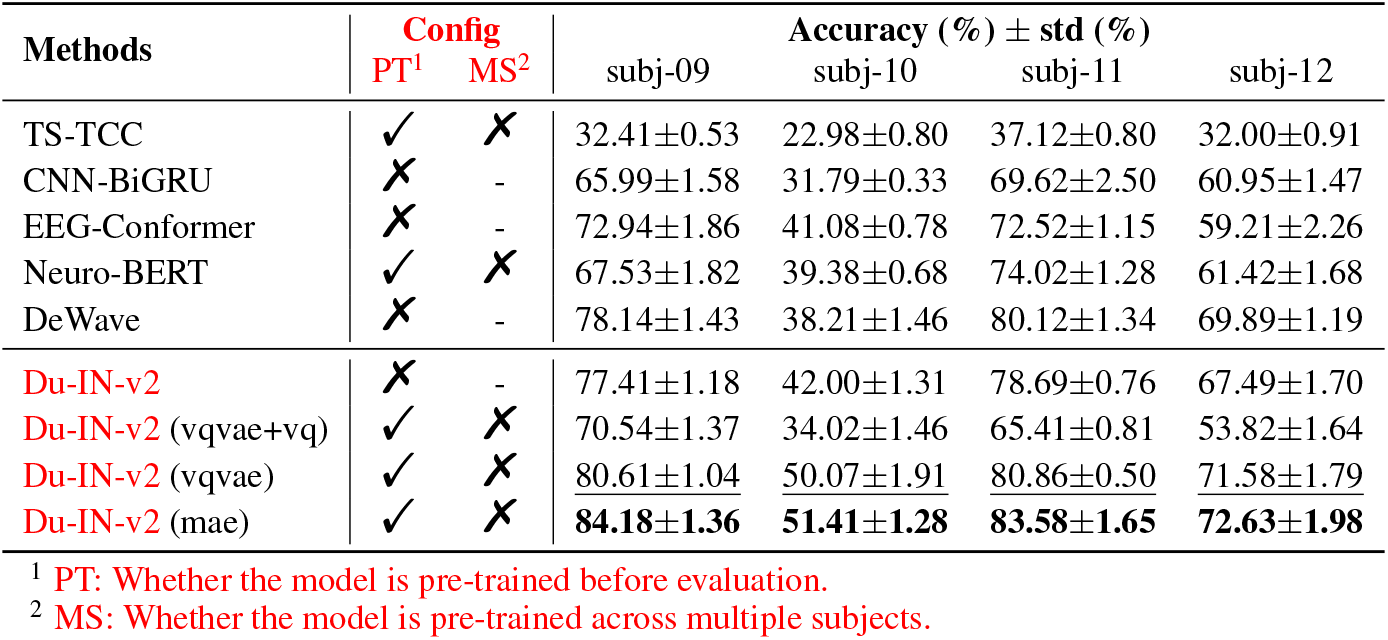
The 49-syllable performance of different methods from subjects (09-12).

**Table 15:**
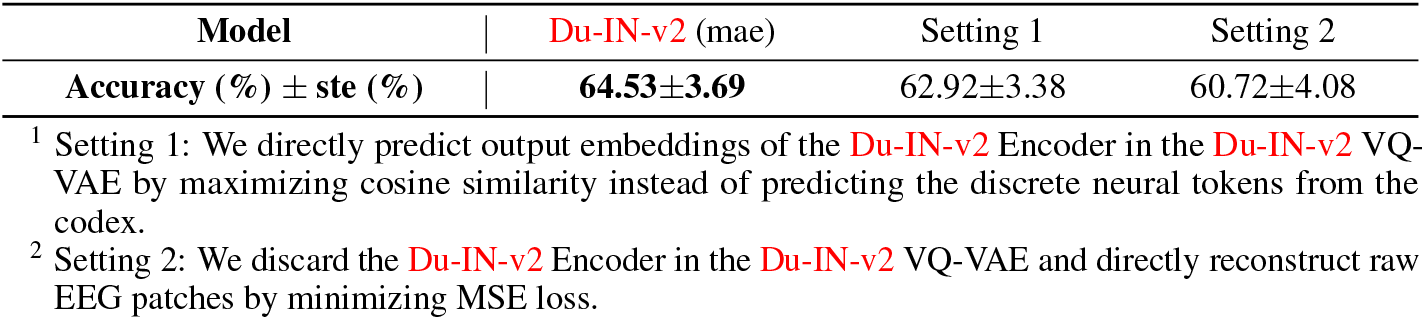
Ablations to validate the effectiveness of vector-quantized neural signal prediction.

**Table 16:**
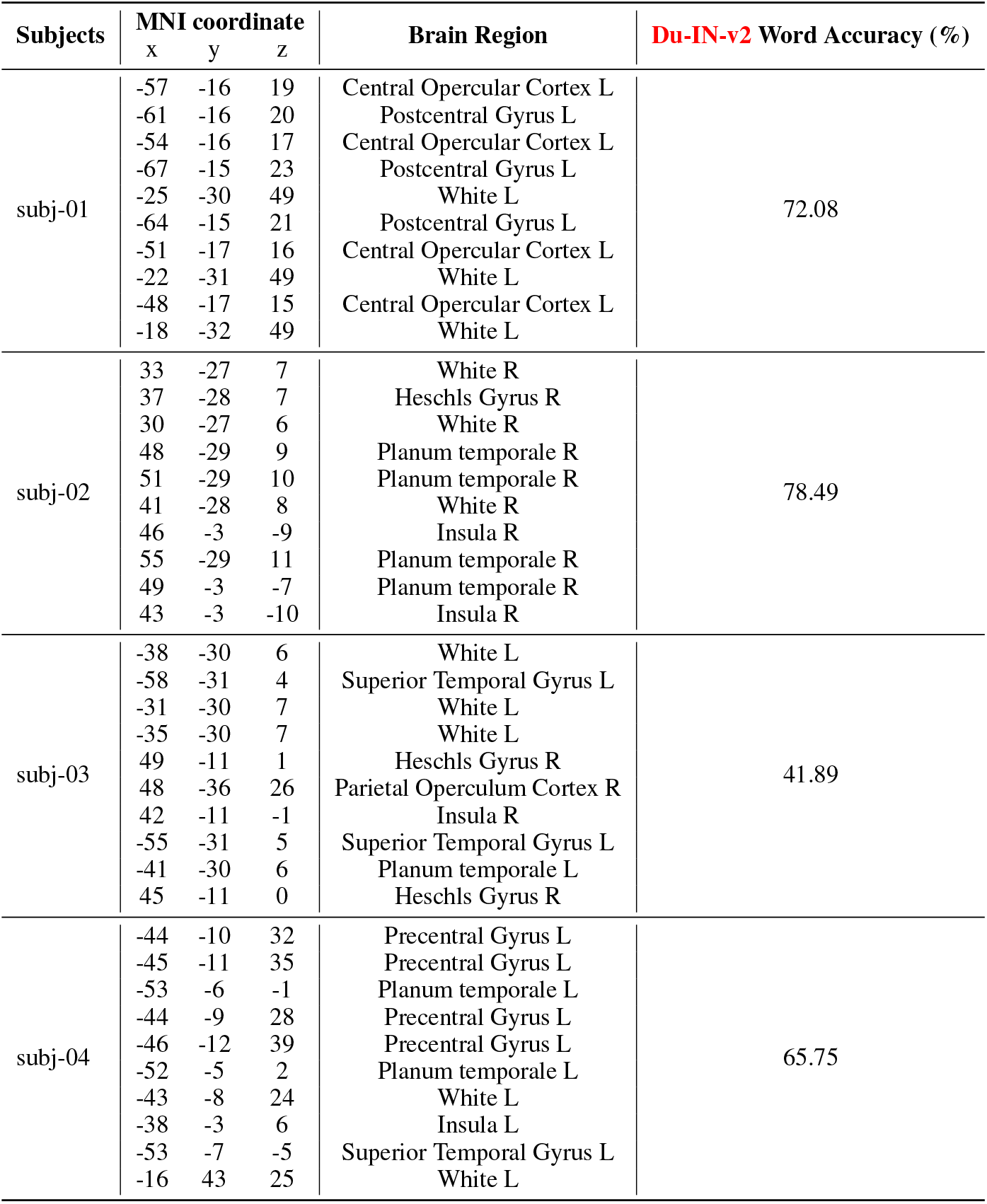
The MNI coordinates and brain region labels of selected channels from subjects (01-04).

**Table 17:**
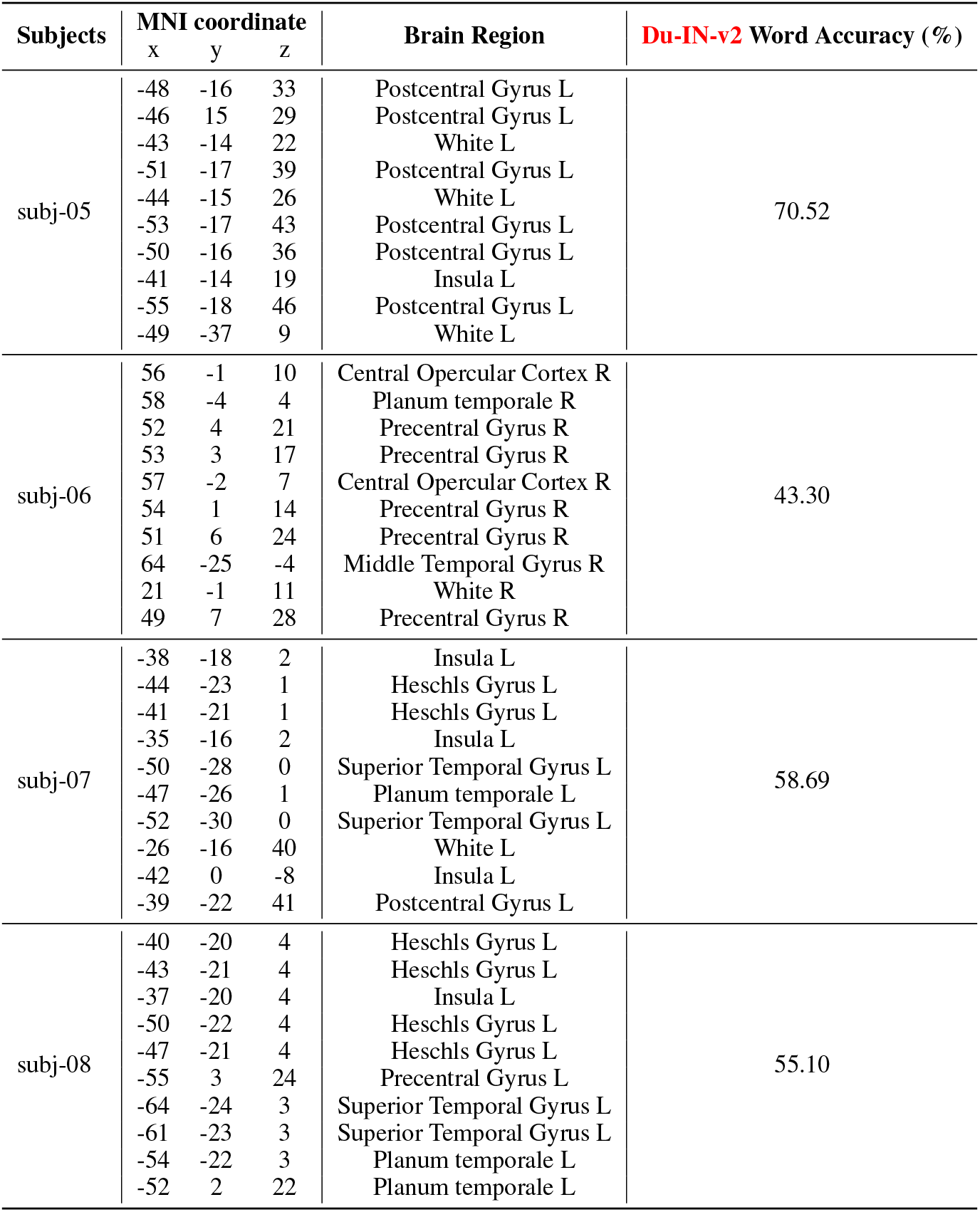
The MNI coordinates and brain region labels of selected channels from subjects (05-08).

**Table 18:**
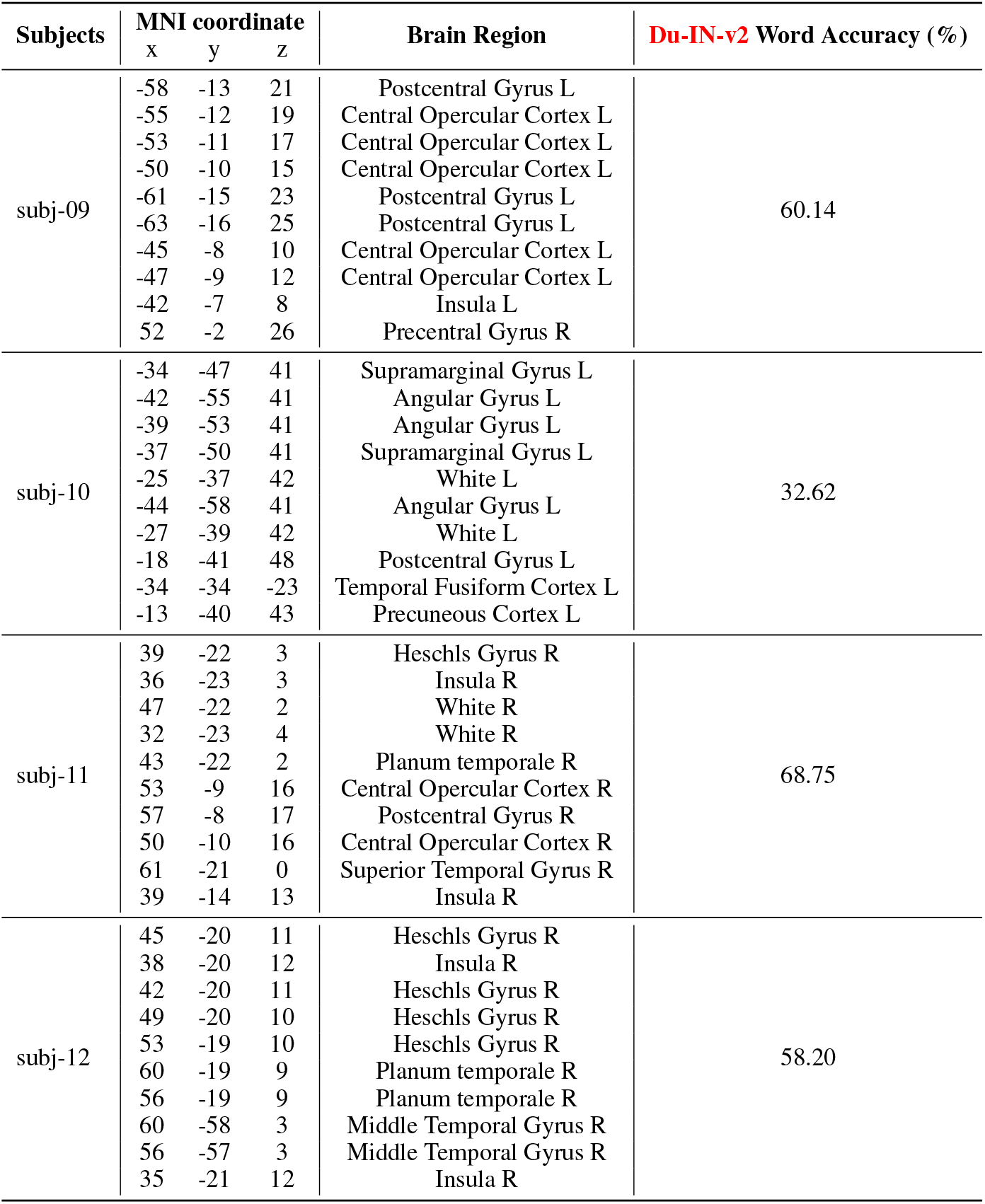
The MNI coordinates and brain region labels of selected channels from subjects (09-12).

#### Discrete Codex

During the Du-IN-v2 VQ-VAE training stage, the Du-IN-v2 VQ-VAE model encodes raw sEEG signals into discrete decoupling codes and then reconstructs the original signal from these codes, as shown in Figure 2. We evaluate performance against varying codex groups (from 0 to 8) to ascertain if the number of codex groups affects the quality of the learned codex. To maintain the capacity of the codex with *G* = 1, we set *N*_*codex*_ to 2048, while for the other settings, *N*_*codex*_ is set to 128. As illustrated in Figure 4 (a), since the channels are pre-selected based on specific brain regions, even a small number of codex groups (e.g., *G* = 4) can effectively decouple different parts of brain dynamics (Chapeton et al., 2022). We also assess performance across different codex sizes (from 32 to 1024) to ascertain if codex size affects the quality of the learned codex. As illustrated in Figure 4 (b), while extremely small codex size lacks representation diversity, extremely large codex size often leads to codex collapse. We suspect that our existing training data might not be adequate for larger codex sizes. Furthermore, our experiments suggest that the model performs best when the dimension of full code z_*q*_(*e*_*i*_), denoted as *G* × *d*_*codex*_ = 4 × 32 = 128, is slightly smaller than the model dimension, *d* = 256, resulting in more effective regularization.

**Figure 4.**
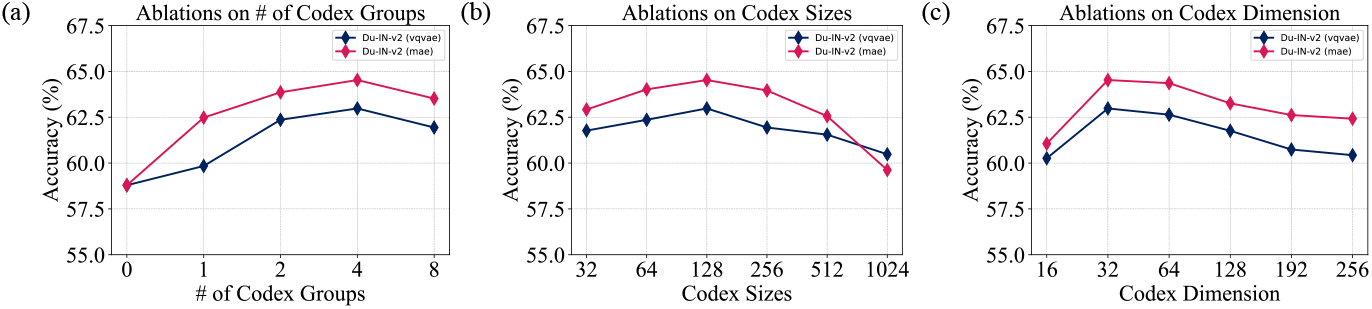
Ablation study on different codex groups, codex sizes, and codex dimensions. For better illustration, we only report the performance difference on the 61-word classification task.

**Figure 5.**
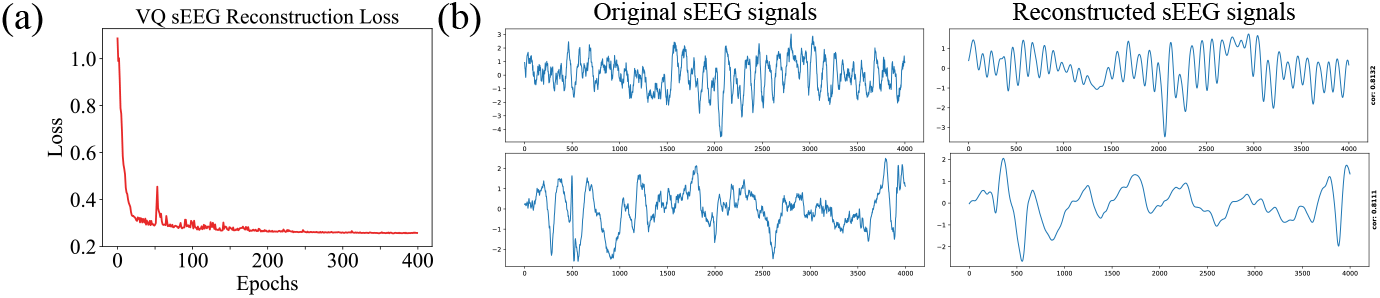
The visualization of Vector-Quantized sEEG Regression. **(a)**.The reconstruction loss curve during the training process of the Du-IN-v2 VQ-VAE model. **(b)**. The visualization of reconstructed sEEG signals.

**Figure 6.**
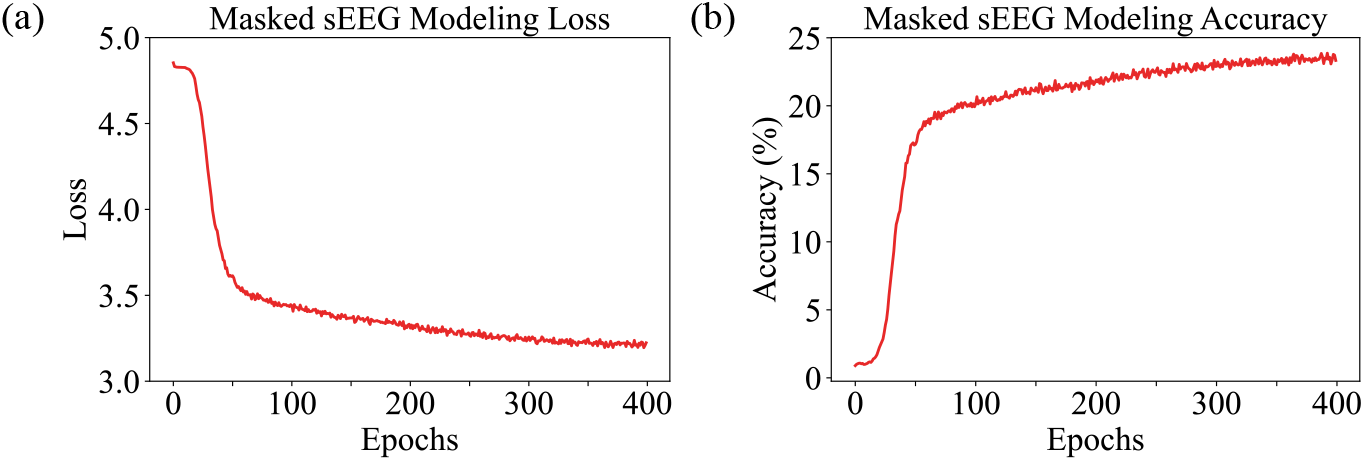
The loss curve and accuracy curve during the training process of the Du-IN-v2 MAE model.

**Figure 7.**
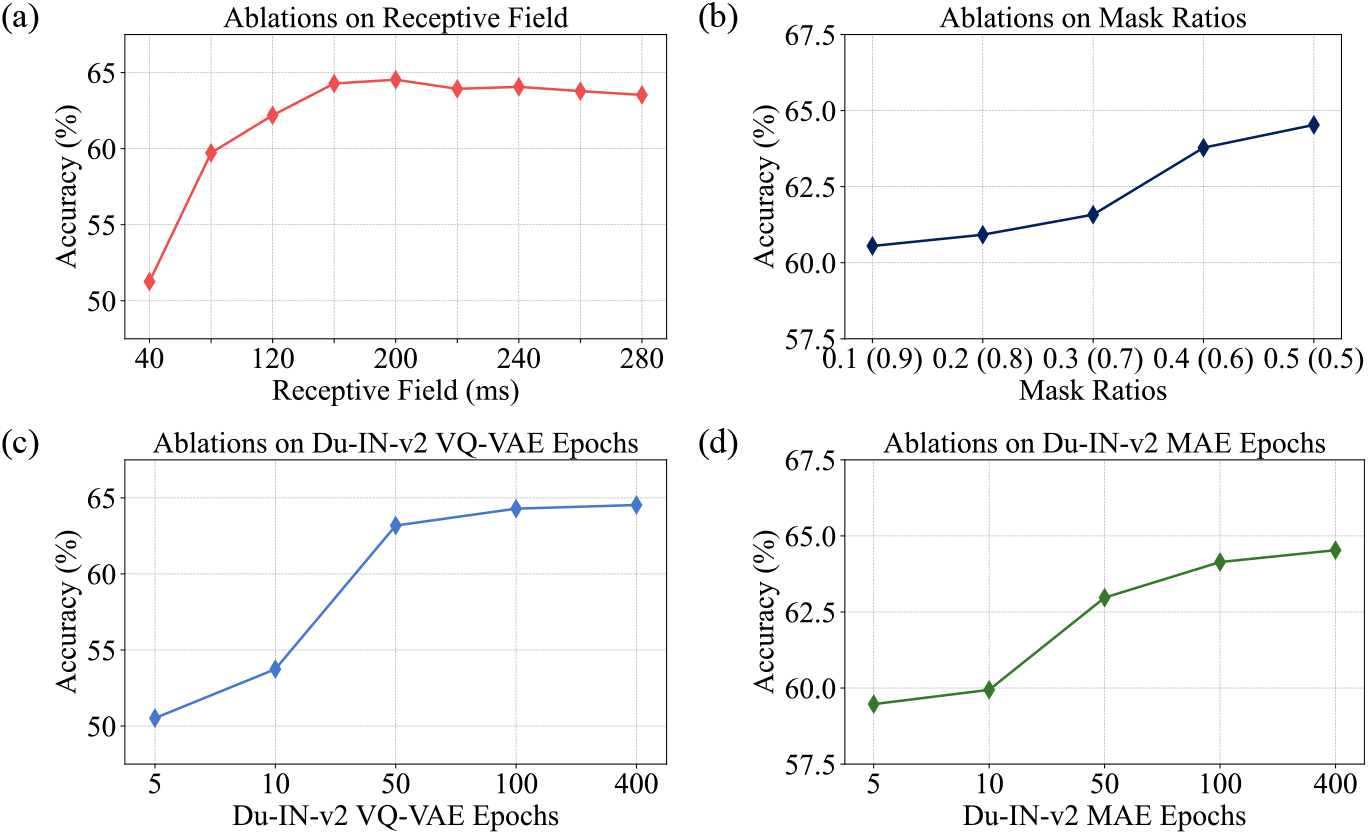
Ablation study on different mask ratios, Du-IN-v2 VQ-VAE epochs, and Du-IN-v2 MAE epochs.

**Figure 8.**
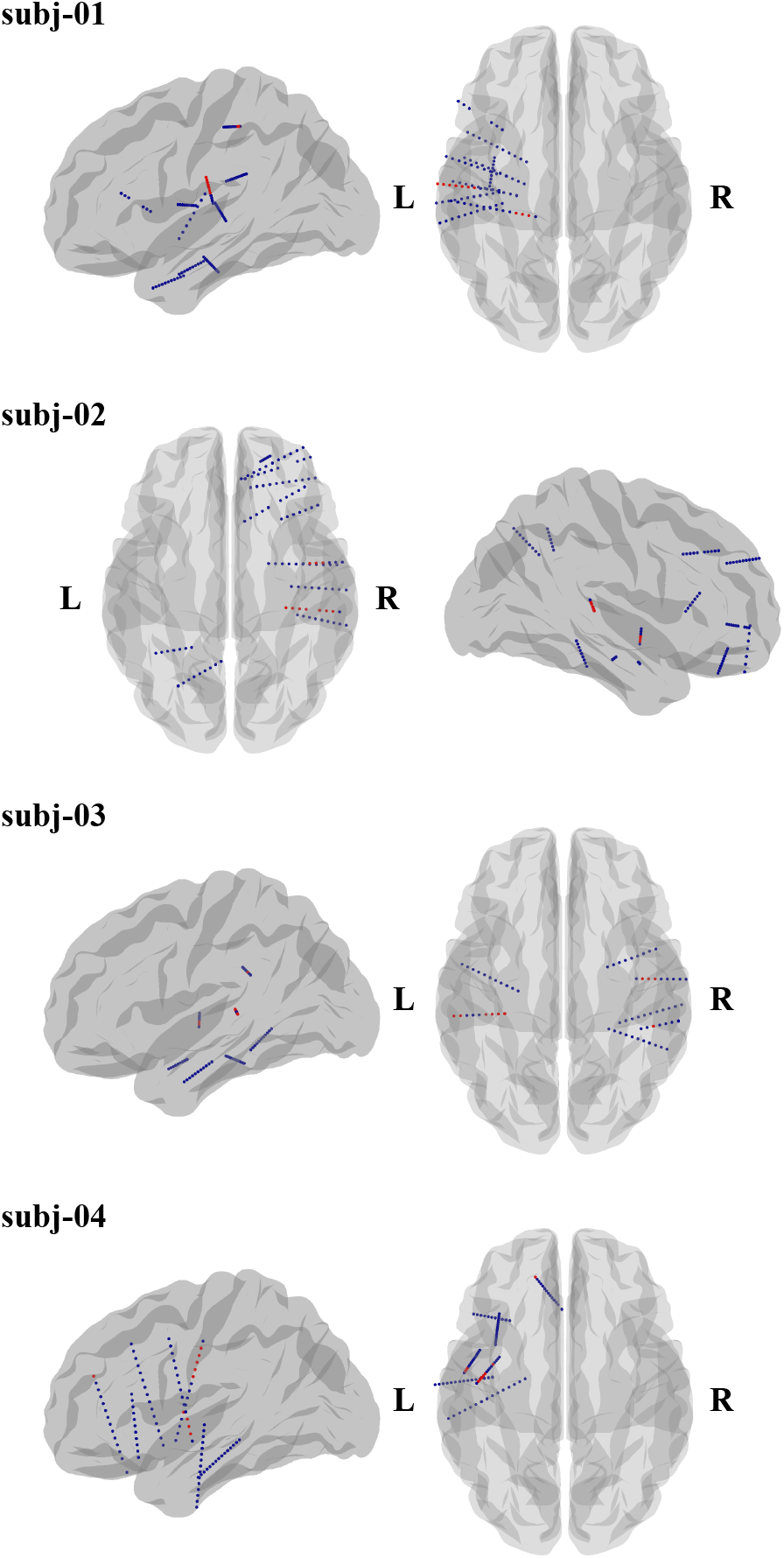
Electrode locations from subjects (01-04).

**Figure 9.**
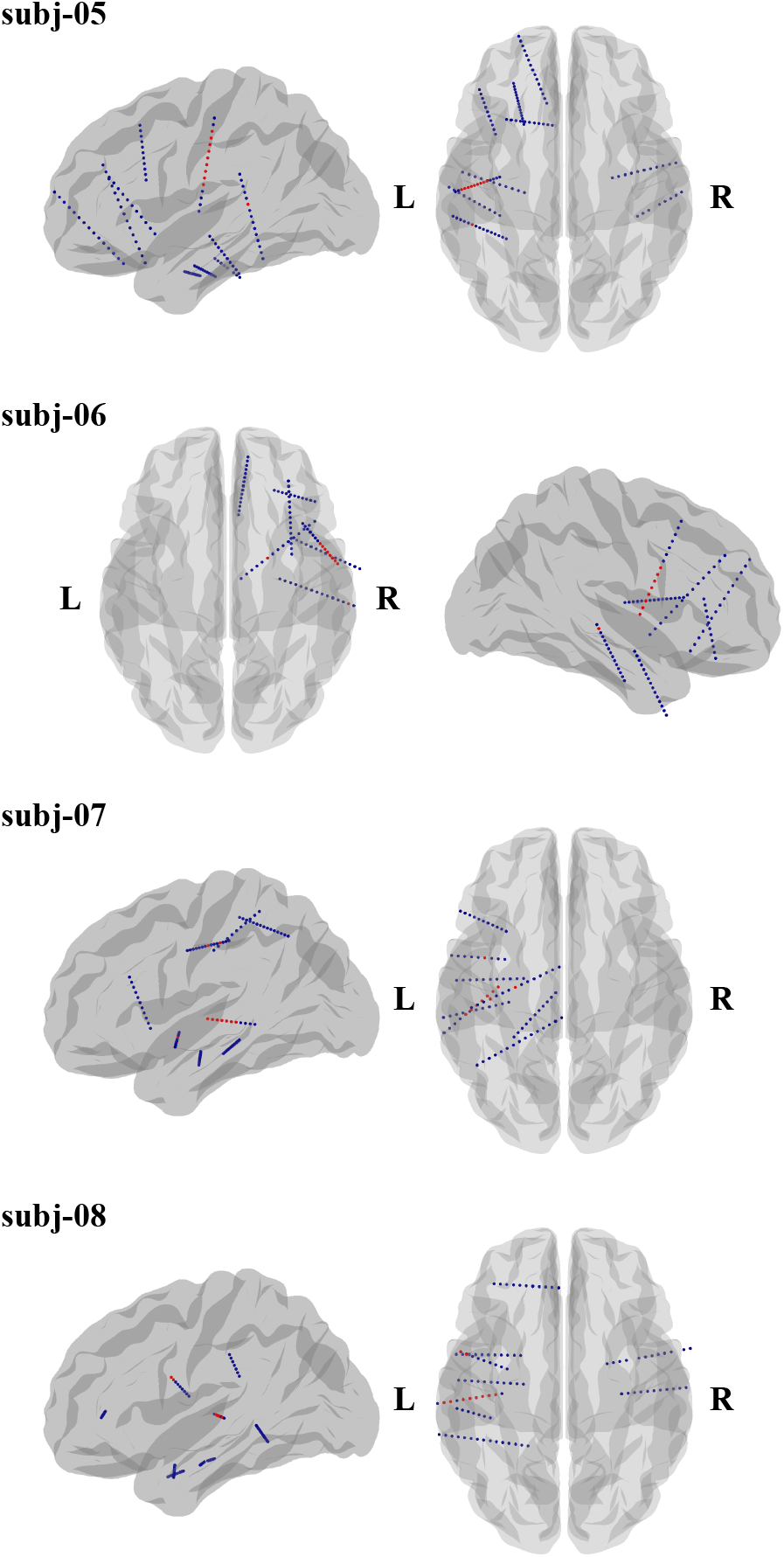
Electrode locations from subjects (05-08).

**Figure 10.**
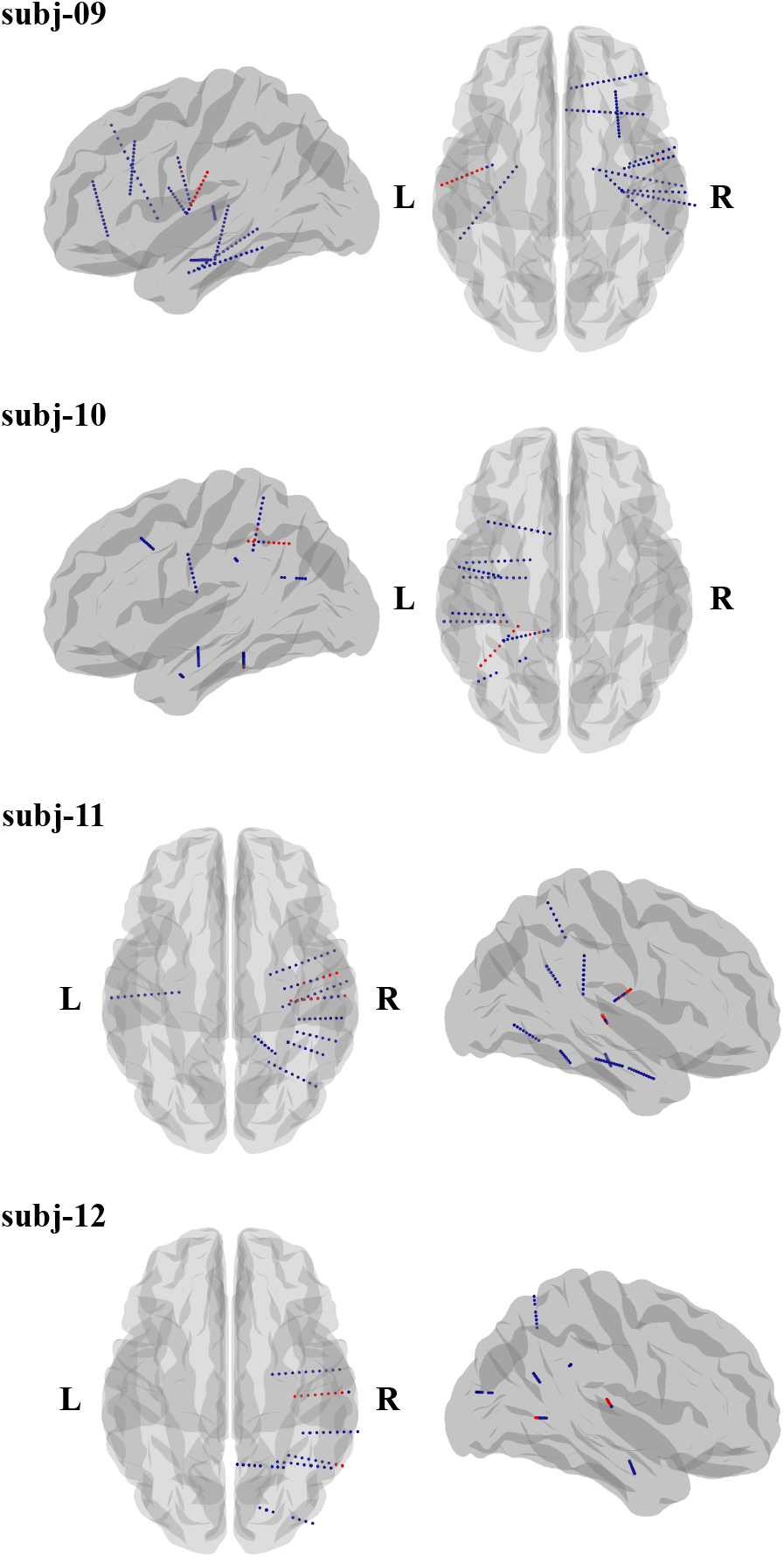
Electrode locations from subjects (09-12).

## 5 CONCLUSION

This paper proposes Du-IN-v2, a general pre-training framework for speech decoding, which learns region-level contextual embeddings through discrete decoupling codex-guided mask modeling on specific brain regions. To evaluate our model, we collect a well-annotated Chinese word-reading sEEG dataset to address the lack of sEEG language dataset. Inspired by neuroscientific findings, we analyze the effective brain regions for speech decoding and achieve the best decoding performance with about one electrode in specific brain regions, which dovetails with the past neuroscientific research on language. Comprehensive experiments demonstrate that our model outperforms the advanced baselines that are designed for either brain signals or general time series, effectively capturing the intricate processing within specific brain regions. It marks a promising neuro-inspired AI approach in BCI. In the end, we hope our work can have implications for future developments in sEEG-based self-supervised models with more consideration over how to build the basic representation units so that the model can maximally benefit from the pre-training stage.

## ETHICS STATEMENT

Experiments that contribute to this work were approved by IRB. All subjects consent to participate. All electrode locations are exclusively dictated by clinical considerations.

Our informed consent signing process is as follows:

1. If the experimental participants are adults and have full civil capacity, we will ask them to sign a written informed consent after the participants have fully informed consent;
2. If the experimental participants are minors or do not have full civil capacity, we will ask the participant’s legal guardian to sign a written informed consent after the participants and their legal guardians have fully informed consent.

Our informed consent form includes the following points:

1. Contact information of research institutions and researchers;
2. Research direction and purpose;
3. Risks involved in the research;
4. Personal information, data, and usage methods to be used in the research;
5. Privacy protection statement (all personal identification information (PII) will not be disclosed);
6. Data storage statement (retained after deleting all personal identification information (PII));
7. Voluntary statement of participants;
8. Statement that participants can withdraw unconditionally at any time.

Our data storage and protection procedures include the following processes:

1. Our data collection, transfer, and analysis tasks are only completed by researchers who have signed relevant confidentiality agreements;
2. The collected raw data will be copied twice as soon as possible, one copy to a storage computer that is not connected to the Internet and encrypted, and the other copy to a mobile hard disk and encrypted and stored offline;
3. The use of the data is only authorized to the research leader and the main researchers (less than 5 people), among which the main researchers can only access data that does not contain personal identification information (PII);
4. After the study is completed, all personal identification information (PII) on both nodes (storage computer, mobile hard disk) will be deleted immediately.

To prevent unauthorized access or possible data leakage, we use double encryption on the storage computer, that is, a static password and a dynamic password (received by mobile phone or email); physical isolation is used on the mobile hard disk, that is, it is locked in a filing cabinet, and the key is only kept by the research leader and the main researchers.

## REPRODUCIBILITY STATEMENT

Code to train models and reproduce the results is submitted as part of the supplementary materials and can be accessed here: TODO, including a demo dataset of 3 subjects for downstream fine-tuning.

## A. DETAILS OF BASELINES

In experiments, we compare our model to the existing supervised or self-supervised methods on brain signals. The details of these baseline models are given here:

- **TS-TCC**(Eldele et al., 2021): A self-supervised model that consists only of a CNN module to capture local features. This model learns robust temporal and discriminative representations from time series by designing a tough cross-view prediction task and a contextual contrasting module. Since sEEG is a unique type of time series, this model is suitable to serve as a baseline for comparison.
- **CNN-BiGRU**(Moses et al., 2021): A supervised model that consists of both CNN module and Bi-GRU module, to capture contextual features from EEG signals. This model is mainly designed for ECoG-based vocal production tasks, similar to ours. Since ECoG and sEEG are both intracranial neural signals of the brain, this model is suitable to serve as a baseline for comparison.
- **EEG-Conformer**(Song et al., 2022): A supervised model that consists of both CNN module and Transformer module, to encapsulate local and global features in a unified EEG classification framework. EEG-Conformer is mainly designed for EEG-based motor imagination tasks. Since the data modes of EEG and sEEG are similar, and the signals primarily pertain to vocal production, this model is suitable to serve as a baseline for comparison.
- **Neuro-BERT**(Wu et al., 2024): A self-supervised model that consists of both CNN module and Transformer module, to encapsulate local and global features. This model learns robust contextual representations from EEG by introducing mask modeling. Since the data modes of EEG and sEEG are similar, this model is suitable to serve as a baseline for comparison.
- **DeWave**(Duan et al., 2023): A supervised model that consists of both Conformer module (Gulati et al., 2020) and Transformer module, to encapsulate local and global features for language decoding. We adopt its encoder, which consists of a 6-layer Conformer and a 6-layer Transformer. Then, we add a classification head, which is also used in our model, for downstream word classification. Since DeWave is also designed for language decoding, this model is suitable to serve as a baseline for comparison.
- **BrainBERT**(Wang et al., 2023): A self-supervised model for sEEG recordings that bridges modern representation learning approaches to neuroscience. BrainBERT builds universal representation based on the superlet spectrograms of one single sEEG channel without modeling the spatial relationships among channels. Since the downstream tasks for BrainBERT are also related to language decoding (e.g., sentence-onset detection, speech vs. non-speech detection, etc.), this model is suitable to serve as a baseline for comparison.
- **Brant**(Zhang et al., 2024): A self-supervised model for sEEG recordings that can capture both long-term temporal dependency and spatial correlation from neural signals. Brant is mainly designed for medicine, serving as a sEEG foundation model. Although Brant mainly evaluates its performance on the low-level modeling tasks (Wu et al., 2022) (e.g., neural signal forecasting, imputation, etc.), Brant achieves SOTA performance on some high-level modeling tasks (e.g., seizure detection). As a foundation model in sEEG pre-training field, this model is suitable to serve as a baseline for comparison.
- **LaBraM**(Jiang et al., 2024): A self-supervised model for EEG recordings that learns generic representations with tremendous EEG data. LaBraM serves as an EEG foundation model, achieving SOTA performance on various downstream EEG tasks. Since the spatial embeddings are pre-defined according to the EEG caps, LaBraM can only be trained within one subject under the sEEG setting. Since the data modes of EEG and sEEG are similar, this model is suitable to serve as a baseline for comparison.
- **LaBraM+PopT**(Jiang et al., 2024; Chau et al., 2024): A self-supervised model based on LaBraM, simply replacing the learnable spatial embeddings with hard-coded spatial embeddings from PopT (Chau et al., 2024) to enable multi-subject pre-training under the sEEG setting.

The detailed implementations of these baseline models are given here:

- For the TS-TCC method (Eldele et al., 2021), the hyper-parameters are optimized for better performance, as they also have different hyper-parameter settings for different datasets in their original implementation. The data samples are resampled to 400Hz. The sizes of convolution kernels are changed to {25, 8, 8} (other attempts include {8, 8, 8}, {15, 8, 8}, {20, 8, 8}, and {30, 8, 8}); the sizes of pooling kernels are changed to {10, 2, 2} (other attempts include {2, 2, 2}, {5, 2, 2}, and {20, 2, 2}); the numbers of pooling strides are changed to {10, 2, 2} (other attempts include {2, 2, 2}, {5, 2, 2}, and {20, 2, 2}). All other hyper-parameters are the same as the original implementation.
- For the CNN-BiGRU method (Moses et al., 2021), the hyper-parameters are the same as the original implementation. The data samples are resampled to the specified sampling rate.
- For the EEG-Conformer method (Song et al., 2022), the hyper-parameters are the same as the original implementation. The data samples are resampled to the specified sampling rate.
- For the Neuro-BERT method (Wu et al., 2024), the hyper-parameters are optimized for better performance, as they also have different hyper-parameter settings for different datasets in their original implementation. The data samples are sampled to 400Hz. The sizes of convolution kernels are changed to {40,} (other attempts include {20,} and {80,}); the numbers of convolution strides are changed to {40,} (other attempts include {20,} and {80,}).
- For the DeWave method (Duan et al., 2023), the hyper-parameters are the same as the original implementation. The data samples are resampled to the specified sampling rate.
- For the BrainBERT method (Wang et al., 2023), the hyper-parameters are optimized for better performance. We change the “nperseg” and “noverlap” arguments of “scipy.signal.stft” function from {400, 350} to {1600, 1400} (other attempts include {200, 175}, {800, 700} and {3200, 2800}).
- For the Brant method (Zhang et al., 2024), the hyper-parameters are optimized based on the original implementation of the Brant-Tiny model. We change the length of the patch segment from 6 seconds to 1 second. We change the linear embedding layer to the convolution embedding layer, which is also used in LaBraM (Jiang et al., 2024). The numbers of convolution filters are {96, 96, 96} (other attempts include {192, 192, 192}); the sizes of convolution kernels are {9, 9, 3} (other attempts include {19, 9, 3} and {9, 9, 3}); the numbers of convolution strides are {5, 5, 1} (other attempts include {10, 5, 1}) and {5, 5, 2}).
- For the LaBraM method (Jiang et al., 2024), the hyper-parameters are the same as the original implementation of the LaBraM-Base model. The data samples are resampled to the specified sampling rate.
- For the LaBraM-PopT method (Jiang et al., 2024; Chau et al., 2024), the hyper-parameters are the same as the original implementation of the LaBraM-Base model. The data samples are resampled to the specified sampling rate.

When evaluating the decoding performance of these baseline models, we follow the same experiment setup of the Du-IN-v2 CLS model:

- For one subject, we split the downstream dataset into training, validation, and testing splits with a size roughly proportional to 80%, 10%, and 10%.
- The data samples are 3 seconds with the specified sampling rate corresponding to each model.
- The samples in the train-set are augmented following the pipeline defined in Appendix C.

For the self-supervised methods, the pre-training setup follows the original setup of each model:

- For the TS-TCC model, we use all sEGG recordings for each subject to pre-train it. The data samples are 4 seconds.
- For the Neuro-BERT model, we use all sEGG recordings for each subject to pre-train it. The data samples are 4 seconds.
- For the BrainBERT model, we use around 180 hours of sEEG recordings from either each subject or 12 subjects for pre-training. This pre-training dataset is larger than the one (approximately 45 hours) used in the original paper. The data samples are 4 seconds.
- For the Brant model, we also use all sEEG recordings from either each subject or 12 subjects to pre-train it. While the total pre-training dataset is smaller than the one (around 2700 hours) used in the original paper, the number of subjects (i.e., the number of sEEG location configurations) is greater than in the original paper. The data samples are 4 seconds.
- For the LaBraM model, we use all sEGG recordings for each subject to pre-train it. The data samples are 4 seconds.
- For the LaBraM-PopT model, we use all sEEG recordings from 12 subjects to pre-train it. The data samples are 4 seconds.

## B. MODEL DETAILS

### B.1. DU-IN-V2 VQ-VAE

The architecture of the Du-IN-v2 VQ-VAE model contains three parts: (1) Du-IN-v2 Encoder, (2) Vector Quantizer, and (3) Du-IN-v2 Decoder. The overall architecture of the “Du-IN-v2 Encoder” is shown in Figure 1. The “Vector Quantizer” is implemented similarly in LaBraM (Jiang et al., 2024). The “Du-IN-v2 Decoder” contains:

- **Transformer Decoder:** A stack of Transformer layers.
- **Time Regression Head:** A stack of 1D Transposed Convolution layers and one linear projection layer.

The hyperparameters for Du-IN-v2 VQ-VAE training are shown in Table 4.

### B.2. DU-IN-V2 MAE

The architecture of the Du-IN-v2 MAE model contains two parts: (1) Du-IN-v2 Encoder, and (2) Token Prediction Head. The overall architecture of the “Du-IN-v2 Encoder” is shown in Figure 1. The hyperparameters of “Du-IN-v2 Encoder” are the same as those in Du-IN-v2 VQ-VAE. It’s worth noting that when training Du-IN-v2 MAE, the weights of the “Du-IN-v2 Encoder” are randomly initialized, instead of loaded from the pre-trained Du-IN-v2 VQ-VAE model. The hyperparameters for Du-IN-v2 MAE training are shown in Table 5.

### B.3. DU-IN-V2 CLS

The Du-IN-v2 CLS model is designed for the 61-word classification task. Similar to Moses et al. (2021), the Du-IN-v2 CLS model aims to decode the corresponding word label *y* from a sequence of raw sEEG signals 𝒳. Specifically, the word label *y* is selected from the pre-determined words in Zheng et al. (2024).

The architecture of the Du-IN-v2 CLS model contains two parts: (1) Du-IN-v2 Encoder, and (2) Label Prediction Head. The overall architecture of the “Du-IN-v2 Encoder” is shown in Figure 1. The hyperparameters of “Du-IN-v2 Encoder” are the same as those in Du-IN-v2 VQ-VAE. It’s worth noting that the “Du-IN-v2 Encoder” weights in Du-IN-v2 CLS can be loaded from either the pre-trained Du-IN-v2 MAE or the pre-trained Du-IN-v2 VQ-VAE. In the ablation experiments shown in Table 2, our models have different suffixes:

- **Du-IN-v2:** The original Du-IN-v2 CLS model. All weights of this model are randomly initialized.
- **Du-IN-v2 (vqvae+vq):** The weights of the “Du-IN-v2 Encoder” in the Du-IN-v2 CLS model are loaded from the pre-trained Du-IN-v2 VQ-VAE. When fine-tuning it on the downstream task, the “Vector Quantizer” in the pre-trained Du-IN-v2 VQ-VAE is inserted between “Du-IN-v2 Encoder” and “Label Prediction Head”. This is the same operation in DeWave (Duan et al., 2023).
- **Du-IN-v2 (vqvae):** The weights of the “Du-IN-v2 Encoder” in the Du-IN-v2 CLS model are loaded from the pre-trained Du-IN-v2 VQ-VAE. This is the same operation in EEGFormer (Chen et al., 2024).
- **Du-IN-v2 (mae):** The weights of the “Du-IN-v2 Encoder” in the Du-IN-v2 CLS model are loaded from the pre-trained Du-IN-v2 MAE. This is the same operation in LaBraM (Jiang et al., 2024).

The “Label Prediction Head” is an MLP with one hidden layer, flattens the output embedding sequence from upstream, and maps this feature embedding to the final prediction through MLP. The hyperparameters for Du-IN-v2 CLS training are shown in Table 6.

### B.4. DU-IN-V2 CTC

The Du-IN-v2 CTC model is designed for the 49-syllable sequence classification task. Similar to Metzger et al. (2023), the Du-IN-v2 CTC model aims to decode the corresponding syllable label sequence *y* from a sequence of raw sEEG signals 𝒳. Considering the difference between English and Chinese, we utilize syllables from Pinyin (Wang, 1973), a widely adopted phonetic representation system based on the Latin alphabet, as basic units that have the potential to support open-set speech decoding tasks. The set of 49 syllables includes:

- 1 CTC blank token (i.e., “-”),
- 1 silence token (i.e., “|”),
- 23 initial syllable tokens,
- 24 final syllable tokens.

Take the pre-determined word “computer” for example, the corresponding syllable label sequence ***y*** is [“|”, “d”, “i”, “an”, “|”, “n”, “ao”, “|”].

The architecture of the Du-IN-v2 CTC model contains two parts: (1) Du-IN-v2 Encoder, and (2) Label Prediction Head. In the ablation experiments shown in Table 2, Du-IN-v2 models with different suffixes are defined the same in Appendix B.3.

The “Label Prediction Head” is an MLP with one hidden layer, flattens the output embeddings with a specified flatten window from upstream, and maps the transformed embedding sequence to the final prediction through MLP. The hyperparameters for Du-IN-v2 CTC training are shown in Table 7.

## C. DATA AUGMENTATION

To enhance the robustness of learned representations during both the pre-training and fine-tuning stages, we apply data augmentation in both datasets.

### Pre-training Dataset

In our implementation, we segment sEEG recordings into 8-second samples with a 4-second overlap. When fetching a sample, we randomly select a starting point between 0 and 4 seconds, then extract a 4-second sample beginning from that point.

### Downstream Dataset

Since a trial lasts for 3 seconds, employing the jittering mentioned above leads to the blending of information from other trials. In our implementation, we segment sEEG recordings into 3-second samples. When fetching a sample, we randomly choose a shift step between 0 and 0.3 seconds, then shift the sample either to the left or right, padding it with zeros.

## D. DU-IN-V2 PRE-TRAINING ANALYSIS

The pre-training of Du-IN-v2 can be interpreted as the training of a variational autoencoder (Kingma & Welling, 2013; Bao et al., 2021). Let *x* denote the original sEEG signal, 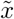 the corrupted sEEG by mask, and *z* the neural tokens. Considering the evidence lower bound (ELBO) of the log-likelihood 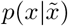, i.e., recovering the original sEEG signal from its corrupted version:

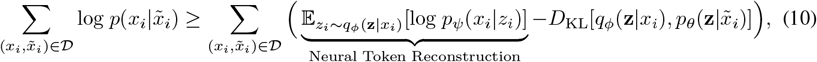

where (1) *q*_*ϕ*_(*z* |*x*) denotes the Du-IN-v2 Encoder in the Du-IN-v2 VQ-VAE that obtains neural tokens; (2) *p*_*ψ*_(*x*| *z*) decodes the original sEEG signal given input neural tokens; (3) 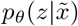 recovers the neural tokens based on the masked sEEG signal, which is our Du-IN-v2 pre-training task.

The whole framework is optimized through a two-stage procedure (Van Den Oord et al., 2017; Razavi et al., 2019). For the first stage, we train the Du-IN-v2 Encoder in the Du-IN-v2 VQ-VAE as a discrete variational autoencoder by minimizing the reconstruction loss 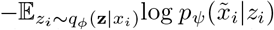 with a uniform prior. For the second stage, we set *q*_*ϕ*_ as well as *p*_*ψ*_ fixed and learn the prior *p*_*θ*_ by minimizing the loss *D*_KL_. For simplicity, *q*_*ϕ*_(**z** |*x*_*i*_) is defined as a one-point distribution with the most likely neural tokens 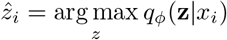. Consequently, we can rewrite Equation 10 as

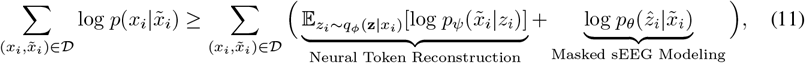

where the first term is the objective for vector-quantized neural signal regression in the first stage (i.e., the Du-IN-v2 VQ-VAE model), and the second term is the objective for Du-IN-v2 pre-training in the second stage (i.e., the Du-IN-v2 MAE model).

## E. VISUALIZATION OF VECTOR-QUANTIZED SEEG REGRESSION

We further visualize how the sEEG signals are reconstructed. As depicted in Figure 5, although some details are missing, the overall trend of the signals is reconstructed well. Meanwhile, there is a stable decrease in the reconstruction loss during training, which indicates the discrete codex does learn high-level information from sEEG signals.

## F. VISUALIZATION OF MASK SEEG MODELING

Figure 6 demonstrates the convergence curves of the total pre-training loss and masked sEEG modeling accuracy of the Du-IN-v2 MAE model. We observe that there is a stable decrease in the mask modeling loss, and the mask modeling accuracy achieves about 20%.

## G. CHANNEL CONTRIBUTION ANALYSIS

For each subject, after training the Du-IN-v2 model (with random initialization) on the downstream dataset, we utilize the weights *W* ∈ ℝ^*C×D*^ of linear projection in the spatial encoder to calculate the contribution scores 𝒮 of channels:

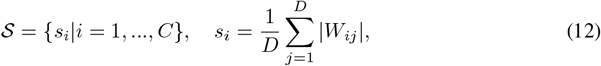

where *C* is the number of channels, *D* is the dimension of projected embedding and | · | gets the absolute value. Then, we normalize 𝒮 using its maximum value to ensure it falls within the [0,1] range. Finally, given the variability in model performance across subjects, we further adjust the channel contribution scores based on the decoding performance of that subject, i.e., 𝒮 = {*s*_*i*_·*p*|*i* = 1, …, *C*}, where *p* represents the decoding performance of that subject.

After calculating the channel contribution scores of all subjects, we project them to the standard brain template according to the MNI (Montreal Neurological Institute) locations of channels, using Nilearn 0.9.2. Since the electrodes are sparsely distributed within the brain, we use Scipy 1.8.1 to interpolate and smooth the channel contribution matrix and use NiLearn to plot the channel contribution map demonstrated in Figure 3 (a).

With the sorted channels within each subject, we evaluate the effect of the number of channels on the decoding performance. For each subject, we evaluate the Du-IN-v2 model with {5, 10, 15, 20, 30, 60} channels (sorted by channel contribution scores), and the averaged performance (across subjects) is demonstrated in Figure 3 (b).

## H EFFECTIVENESS OF REGION-SPECIFIC CHANNEL SELECTION

DeWave (Duan et al., 2023) successfully reconstructs 128-channel EEG signals with the setting of vector-quantizer (*G* = 1; *N*_*codex*_ = 2048). However, this is not the case under the sEEG setting, which is shown in Table 8. We should note that sEEG signals are fundamentally different from EEG signals due to (1) the high information density and (2) the high specificity of different regions. Due to the desynchronization nature (Buzsaki, 2006) of the brain during awake tasks, only specific brain regions are related to tasks. Therefore, only after region-specific channel selection, the Du-IN-v2 VQ-VAE model can successfully reconstruct the original signals, thus identifying the fine-grained state of brain regions.

## I. SUBJECT-WISE EVALUATION

### I.1 61-WORD CLASSIFICATION TASK

The detailed performance of different methods from each subject is provided in Table 9, Table 10, and Table 11, with the best in **bold** and the second underlined. For model comparison, we report the average and standard deviation values (within each subject) on six different random seeds to obtain comparable results. “std” means standard deviation.

### I.2 49-SYLLABLE SEQUENCE CLASSIFICATION TASK

The detailed performance of different methods from each subject is provided in Table 12, Table 13, and Table 14, with the best in **bold** and the second underlined. For model comparison, we report the average and standard deviation values (within each subject) on six different random seeds to obtain comparable results. “std” means standard deviation.

## J. EFFECTIVENESS OF VECTOR-QUANTIZED NEURAL SIGNAL PREDICTION

To verify the effectiveness of vector-quantized neural signal prediction, we elaborate on two types of experimental settings as illustrated in Table 15. The comparison between Du-IN-v2 and Setting 1 demonstrates that the codex is effective for masked sEEG modeling. The comparison between Du-IN-v2 and Setting 2 demonstrates that introducing the codex can prevent the model from focusing too much on reconstructing details, thus enabling the Du-IN-v2 MAE to learn better contextual embeddings.

## K. ADDITIONAL ABLATION STUDY

### K.1 ABLATION ON PERCEPTION TIME WINDOW

We conduct the ablation study on the model structure for the spatial encoder described in Section 3.2. As the spatial encoder transforms the sEEG patches with overlap into tokens, it compresses the sEEG signals for perception. We conduct an ablation study of different receptive fields and report it in Figure 7 (a). The model performance notably drops with a receptive field smaller than 80ms and gradually declines as the receptive field exceeds 240ms. The model reaches a small peak around 160ms to 240ms. We think this phenomenon is rational since sEEG is known for its ability to capture the rapid dynamics of specific brain regions precisely.

### K.2.ABLATION ON MASK RATIO

In this experiment, we conduct different mask ratio settings to explore its impact. It is noted that we introduce the symmetric masking strategy, so we only need to validate half of the mask ratios. As the mask ratio is set to *r*, the symmetric masking will mask the 1 − *r* proportion of sEEG tokens. Ablation results in Figure 7 (b) show the optimal mask ratio for our dataset is 0.5 (0.5).

### K.3. ABLATION ON PRE-TRAINING EPOCHS

The impact of the number of pre-training epochs (of the Du-IN-v2 VQ-VAE model) is demonstrated in Figure 7 (c). We use the checkpoints of the specified epochs to pre-train the Du-IN-v2 MAE model for 400 epochs. Once the reconstruction loss of the Du-IN-v2 VQ-VAE model converges, the Du-IN-v2 VQ-VAE model can extract the state of the brain region well, thus leading to better performance.

The impact of the number of pre-training epochs (of the Du-IN-v2 MAE model) is demonstrated in Figure 7 (d). We use the checkpoints of the specified epochs for downstream classification. Once the mask modeling loss of the Du-IN-v2 MAE model converges, the Du-IN-v2 MAE model learns robust contextual embeddings, thus leading to better performance.

## L. LIMITATIONS

Despite Du-IN-v2’s enhancements in speech decoding via discrete codex-guided mask modeling, it is still restricted to close-set speech decoding tasks (i.e., the word set only includes 61 pre-determined words). However, a parallel to our work (Feng et al., 2023), which follows previous works (Herff et al., 2015; Soroush et al., 2023), builds an acoustic-inspired model that can decode arbitrary Chinese words by predicting syllable components (initials, finals, tones). Although their method requires a large amount of labeled data, their experimental design mirrors ours closely. The difference lies in the requirement for the subject to repeat syllable components instead of entire words. Therefore, with slight modifications, our model can support open-set speech decoding tasks. Besides, we also provide the result of the 49-syllable sequence classification task following Metzger et al. (2023), which shows the potential generality of our approach on open-set decoding tasks.

Additionally, the experiments in this paper are restricted to the vocal production part of language decoding, i.e. speech decoding. A more interesting but difficult task is to decode language from the semantic level, in which large language models have been wildly used to improve the model performance (Tang et al., 2023; Duan et al., 2023). However, due to the locality of sEEG recordings, it is still under exploration whether sEEG recordings can fully capture semantic-related information across brain regions.

## M. SUBJECT-WISE ELECTRODE LOCATIONS

We provide detailed information on the locations of the implanted sEEG electrodes for each subject. Red channels are the top 10 channels (selected through channel contribution analysis) for both pre-training and downstream evaluation, as described in Section 4.3. As the majority of subjects have sEEG electrodes implanted on only one side of their brains to locate the source of epilepsy, we provide side views of either the left or right brain areas here.

## N. SUBJECT-WISE SELECTED CHANNELS

The MNI coordinates and brain region labels (according to Harvard-Oxford cortical and subcortical structural atlases Desikan et al. (2006)) for selected channels are listed below. The channels for each subject are arranged in descending order based on their contribution scores.

## Footnotes

1 The term "group-level" includes "brain-level" and "region-level," and is distinct from "channel-level."

